# A universal influenza mRNA vaccine candidate boosts T-cell responses and reduces zoonotic influenza virus disease in ferrets

**DOI:** 10.1101/2022.08.02.502529

**Authors:** Koen van de Ven, Josien Lanfermeijer, Harry van Dijken, Hiromi Muramatsu, Caroline Vilas Boas de Melo, Stefanie Lenz, Florence Peters, Mitchell B Beattie, Paulo J C Lin, José A. Ferreira, Judith van den Brand, Debbie van Baarle, Norbert Pardi, Jørgen de Jonge

**Author notes:** correspondence should be sent to and.

## Abstract

Universal influenza vaccines have the potential to protect against continuously evolving and newly emerging influenza viruses. T cells may be an essential target of such vaccines as they can clear infected cells through recognition of conserved influenza virus epitopes. We evaluated a novel T cell-inducing nucleoside-modified mRNA vaccine that encodes the conserved nucleoprotein, matrix protein 1 and polymerase basic protein 1 of an H1N1 influenza virus. To mimic the human situation, we applied the mRNA vaccine as a prime-boost regimen in naïve ferrets (mimicking young children) and as a booster in influenza-experienced ferrets (mimicking adults). The vaccine induced and boosted broadly-reactive T cells in the circulation, bone marrow and respiratory tract. Booster vaccination enhanced protection against heterosubtypic infection with potential pandemic H7N9 influenza virus in influenza-experienced ferrets. Our findings show that mRNA vaccines encoding internal influenza virus proteins are a promising strategy to induce broadly-protective T-cell immunity against influenza viruses.

## Introduction

Influenza viruses infect 5-15% of the world population annually, resulting in approximately 290-650 thousands of deaths worldwide [1, 2]. While vaccines mitigate influenza virus-induced morbidity and mortality, the effectiveness of inactivated influenza virus vaccines is insufficient [3-5]. These vaccines mainly induce strain-specific immunity and are therefore limited in their ability to protect against mutated or newly introduced influenza virus strains. Animal-to-human transmissions of influenza A viruses pose a particular risk, as seasonal influenza vaccination does not offer protection against these strains. There are ample examples of influenza viruses crossing the species barrier and causing a pandemic, with the Spanish flu of 1918 as the most dramatic known example [6, 7]. Recent zoonotic transmissions of highly pathogenic avian influenza virus – like H5N1 and H7N9 – have occurred frequently and are associated with high mortality rates [8, 9]. Especially alarming is the recent rise in outbreaks of these viruses on poultry farms and among migrating birds in Europe and other parts of the world [10]. Although human-to-human transmission of these viruses has been limited so far, experimental work indicates that only a few mutations are required to enhance transmission among humans, highlighting their pandemic potential [11-13]. This emphasizes the ongoing threat posed by influenza viruses and the requirement for a broadly-reactive influenza vaccine that protects against all influenza subtypes.

The narrow protection of inactivated influenza virus vaccines is mainly due to the induction of strain-specific antibodies against the highly variable globular head domain of influenza virus hemagglutinin (HA) [14]. New vaccine concepts strive to provide a wider range of protection by inducing responses against more conserved protein domains [15]. One way to achieve this is by inducing T-cell responses, as they can recognize epitopes derived from conserved influenza proteins such as nucleoprotein (NP), matrix protein 1 (M1) and polymerase basic protein 1 (PB1) [16-18]. T-cells can clear infected cells and T-cell immunity is associated with improved influenza disease outcome in humans [19-23]. In addition, animal models have confirmed that T cells can protect against heterosubtypic influenza virus infections [24-30]. For these reasons, various new influenza vaccine concepts focus on inducing protective T-cell immunity [14].

In recent years, lipid nanoparticle (LNP)-encapsulated nucleoside-modified mRNA (mRNA-LNP) has shown to be a potent novel vaccine format against influenza and other infectious diseases [31, 32]. The potency of the mRNA-LNP platform has been demonstrated by the rapid development and successful world-wide use of mRNA-LNP-based SARS-CoV-2 vaccines [33]. mRNA-LNP induces both T-cell and antibody responses [34-38] and is therefore an interesting platform for novel influenza vaccines. Additionally, mRNA-LNP vaccines can be rapidly produced and are easily adjusted to new emerging viral variants [39]. Multiple influenza vaccines based on mRNA-LNP are currently in development, with promising early results [40-43]. These vaccines, however, primarily focus on inducing humoral responses against HA, without utilizing the potential of T-cell immunity against conserved internal influenza proteins.

There is still very limited information about the potential of mRNA-LNP vaccines for inducing broadly-protective T-cell responses against influenza virus infections. We set out to remedy this knowledge-gap by evaluating the immunogenicity and protective efficacy of a novel mRNA-LNP influenza vaccine in a highly relevant ferret model. We have previously shown in ferrets that circulating and respiratory T cells recognize conserved influenza virus epitopes and can protect against heterosubtypic influenza virus infection [25]. Here, we investigated if we could induce and enhance this protective immunity by vaccination with nucleoside-modified mRNA-LNP encoding for three conserved internal proteins of H1N1 influenza virus, NP, M1 and PB1 (mRNA-Flu). To mimic the human situation – which consists of both naïve young children and influenza-experienced individuals – we evaluated mRNA-Flu as a prime-boost regimen in naïve ferrets (a model for naïve children) and as a booster in influenza-experienced ferrets (a model for influenza-experienced individuals). Both strategies successfully induced and boosted systemic and respiratory T-cell responses, but mRNA-Flu vaccination in influenza-experienced ferrets resulted in higher and broader responses. Moreover, mRNA-Flu booster immunization reduced disease severity in influenza-experienced ferrets after challenge with a potential pandemic avian H7N9 influenza virus, whereas mock-boosted influenza-experienced ferrets were not protected. Our results demonstrate that broadly-reactive T-cell immunity is boosted by a nucleoside-modified mRNA-LNP vaccine that encodes several internal influenza virus proteins. This mRNA-LNP vaccine enhanced protection against heterosubtypic influenza infection and is a promising strategy for the development of a universal influenza vaccine.

## Results

### Study set-up

We designed the mRNA vaccine based on the NP, M1, and PB1 proteins of H1N1 influenza virus (mRNA-Flu) since these proteins are highly conserved (Supplemental Table 1) and immunogenic in humans [25, 44]. To model mRNA-Flu vaccination in both naïve and influenza-experienced humans, we followed a prime-boost strategy with different regimens (Fig. 1a). Naïve ferrets were prime-boosted by intramuscular (i.m.) mRNA-Flu vaccination on days 0 and 42, modelling naïve individuals (group mRNA/mRNA). Another group of ferrets was primed on day 0 by intranasal (i.n.) infection with 10^6^ TCID_50_ A/California/07/2009 (H1N1) influenza virus followed by booster vaccination with mRNA-Flu on day 42 to mimic vaccination of influenza virus-experienced individuals (group H1N1/mRNA). As a control for this treatment, another group of ferrets received the same priming (H1N1 infection), but a mock booster with mRNA-LNP encoding for firefly luciferase (group H1N1/mock) on day 42. A placebo group that received only phosphate-buffered saline (PBS) as a prime-boost served as a negative control. The positive control consisted of ferrets that were primed by H1N1 infection and boosted with 10^6^ TCID_50_ A/Uruguay/217/2007 (H3N2) influenza virus, as a secondary heterosubtypic influenza infection is a very potent booster of T-cell responses (group H1N1/H3N2) [25]. Blood was collected at 0, 14, 42, 56 and 70 days post priming (dpp). Four weeks after the booster (70 dpp), ferrets were euthanized to study systemic and local T-cell responses.

**Figure 1:**
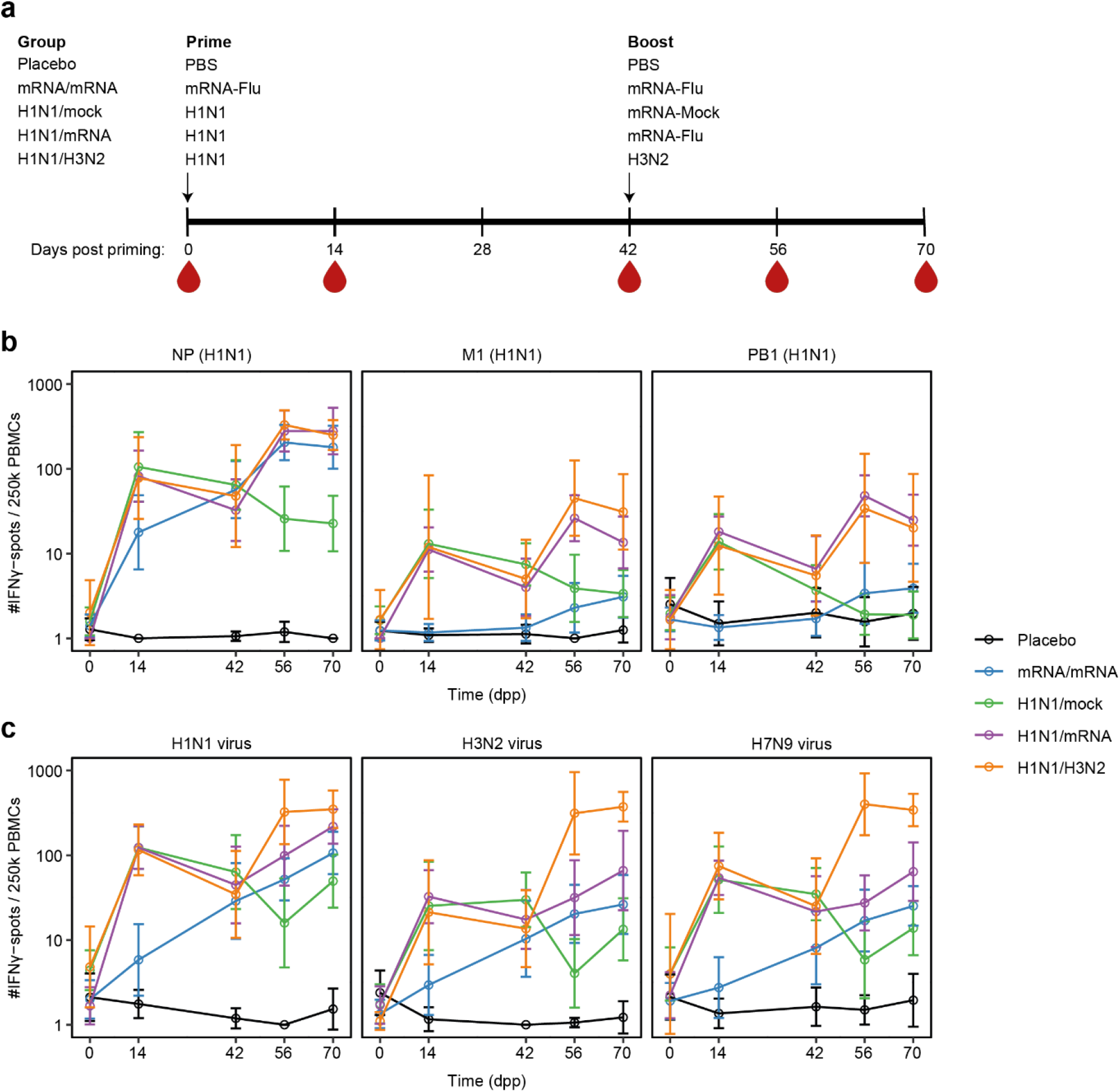
Cellular responses in blood after prime-boost immunization with mRNA-Flu. **a)** Study layout depicting the prime-boost strategy. On day 0, ferrets were primed intranasally with PBS, 10^6^ TCID_50_ H1N1 influenza virus (A/California/07/2009) or primed intramuscularly with mRNA-LNPs encoding for NP, M1 and PB1 (50 μg per mRNA-LNP; mRNA-Flu). Ferrets primed with PBS (group placebo) or mRNA-Flu (group mRNA/mRNA) received the same treatment as booster 42 days post priming (dpp). H1N1-primed ferrets were boosted intramuscularly with mRNA-Flu (H1N1/mRNA-Flu), mRNA-LNP encoding firefly luciferase (50 μg; H1N1/mock) or boosted intranasally with 10^6^ TCID_50_ H3N2 influenza virus (H1N1/H3N2; A/Uruguay/217/2007). Blood was collected on 0, 14, 42, 56 and 70 dpp. Ferrets were euthanized 70 dpp to study cellular responses in tissues. **b, c)** Cellular responses measured by IFNγ ELISpot after 20 hours stimulation of PBMCs with **b)** H1N1 NP, M1 and PB1 overlapping peptide pools or **c)** live influenza viruses H1N1, H3N2 or H7N9 (A/Anhui/1/2013). Data were corrected for medium background and are visualized as geometric mean + geometric standard deviation. n = 7 for H1N1/H3N2 and n = 12-14 for all other groups. Statistics are detailed in Supplemental data file 1.

### An mRNA-based T-cell vaccine induces and boosts systemic cellular responses against conserved influenza virus proteins

We evaluated the cellular responses induced by mRNA-Flu vaccination by stimulation of peripheral blood mononuclear cells (PBMCs) from immunized ferrets with overlapping peptide pools of H1N1 NP, M1 and PB1 in IFNγ ELISpot assays. A single dose of mRNA-Flu induced cellular responses against NP, but not to M1 and PB1 at 14 dpp (Fig. 1b and Supplemental Fig. 1a). Responses were stronger and broader in H1N1 influenza virus-primed ferrets as they displayed responses against NP, M1 and PB1. The cellular response against NP in mRNA-primed ferrets increased further between 14 and 42 dpp, while this response was already contracting in H1N1-primed ferrets. This might be due to the long availability of influenza antigens produced from the mRNA-LNP vaccines after i.m. immunization [45].

mRNA-Flu vaccination at 42 dpp boosted existing cellular responses, irrespective of whether ferrets were initially primed with mRNA-Flu or H1N1 influenza (Fig. 1b). At 56 and 70 dpp, NP-specific responses were similar between mRNA/mRNA and H1N1/mRNA ferrets. Responses against M1 and PB1 were still weaker in the mRNA/mRNA group, although they were clearly boosted as approximately half of the animals developed cellular responses after the second vaccination (Fig. 1b and Supplemental Fig. 1b). Importantly, NP-specific cellular responses in mRNA/mRNA and H1N1/mRNA ferrets were similarly robust to that measured in H1N1-experienced ferrets boosted with H3N2 influenza virus infection. This finding indicates that nucleoside-modified mRNA-LNP vaccination can be as effective in boosting existing T-cell responses as a heterosubtypic influenza infection.

Based on the high level of protein conservation of internal influenza virus proteins (>90%; Supplemental Table 1), T cells induced by mRNA-Flu or H1N1-priming should respond against a wide range of influenza viruses. Indeed, cellular responses measured in PBMCs after stimulation with H1N1 peptide pools correlated strongly with responses obtained with peptide pools specific for H2N2 influenza virus (A/Leningrad/134/17/57; Supplemental Fig. 1c). Live virus stimulations confirmed these findings as we observed substantial responses against heterosubtypic influenza viruses H3N2, H5N1 (A/Vietnam/1204/2004) and H7N9 (A/Anhui/1/2013) (Fig. 1c and Supplemental Fig. 1d). In conclusion, immunization with mRNA-Flu induces and boosts a cellular response that is cross-reactive with a wide range of influenza viruses due to targeting conserved influenza virus epitopes.

### The mRNA-based T-cell vaccine induces and boosts cellular responses in the respiratory tract and bone marrow

T cells located in the respiratory tract are essential for protection against heterosubtypic influenza virus infections [28, 46]. To determine if mRNA-Flu vaccination is also able to induce and boost T-cell responses in the respiratory tract, we assessed cellular immune responses in the bronchoalveolar lavage (BAL) fluid and nasal turbinates (NT) of immunized ferrets by IFNγ ELISpot at 70 dpp. Despite i.m. administration, mRNA-Flu induced robust cellular responses against NP in the NT, but not in the BAL fluid of mRNA/mRNA ferrets (Fig. 2a, supplemental Fig. 1d). The effect of mRNA-Flu vaccination was even more potent in H1N1-primed ferrets. Vaccination effectively increased NP-, M1- and PB1-specific T-cell responses in the NT of H1N1/mRNA ferrets relative to H1N1/mock and mRNA/mRNA ferrets. NP-responses in the BAL fluid of H1N1/mRNA ferrets also demonstrated an increase compared to H1N1/mock ferrets. Responses against homologous (H1N1) and heterosubtypic (H3N2, H5N1, H7N9) influenza viruses were also higher in the NT (significant) and BAL (trend) of H1N1/mRNA ferrets compared to mRNA/mRNA and H1N1/mock ferrets. All groups that were initially primed intranasally with H1N1 influenza virus displayed stronger cellular responses in the NT than the mRNA/mRNA group, irrespective of whether they received a booster, suggesting that the site of priming dictates the response.

**Figure 2:**
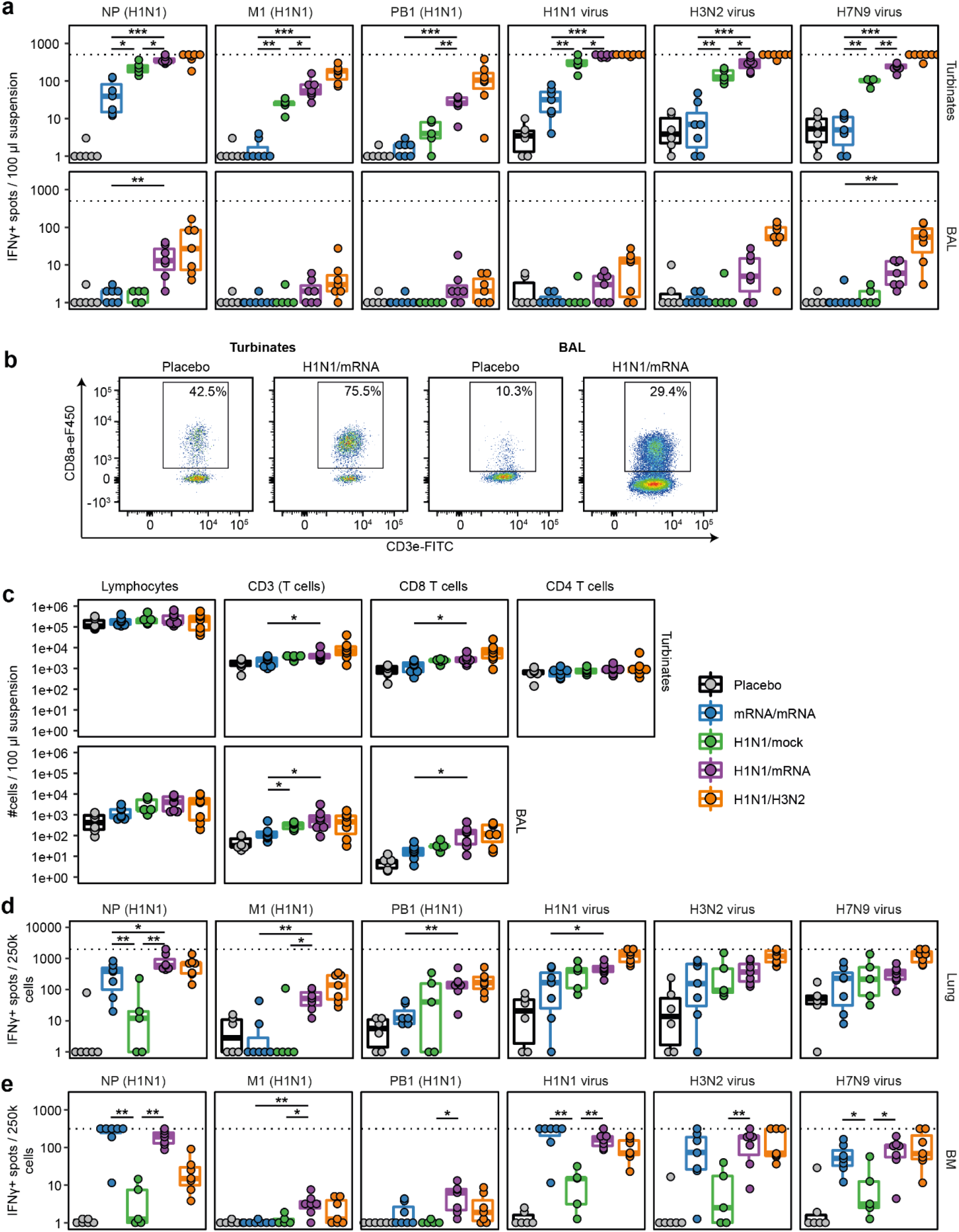
Cellular responses and counts in respiratory compartments and bone marrow of immunized ferrets. **a)** Cellular responses measured by IFNγ ELISpot after 20 hours stimulation with overlapping H1N1 peptide pools or live influenza virus using cells derived from nasal turbinates and bronchoalveolar lavage (BAL) fluid. **b, c)** Cell counts in nasal turbinates and BAL as measured by flow cytometry. **b)** FACS plot displaying the CD8^+^ T cell population in representative turbinate and BAL samples. **c)** Count of different cell populations per 100 μl of suspension. CD4^+^ T cell counts are not displayed for BAL as the αCD4-APC staining was not consistent between BAL samples. **d, e)** Cellular responses measured by IFNγ ELISpot after 20 hours stimulation with overlapping H1N1 peptide pools or live influenza virus of cells derived from **d**) lung or **e**) bone marrow (BM). ELISpot data were corrected for medium background. Boxplots depict the median, 25% and 75% percentile, where the upper and lower whiskers extend to the smallest and largest value respectively within 1.5* the inter quartile ranges. In panels a and c-e, each dot represents one animal and n = 5-7. For visualization purposes, only comparisons between groups mRNA/mRNA, H1N1/mock and H1N1/mRNA are shown. An overview of all statistical comparisons is detailed in Supplemental data file 1. * = p < 0.05, ** = p < 0.01, *** = p < 0.001.

To determine whether mRNA-Flu vaccination also increased absolute T-cell numbers in the respiratory tract, we measured cell counts in the NT and BAL by flow cytometry. Compared to placebo ferrets, T-cell counts (CD3^+^) in the NT were only significantly increased in H1N1/mRNA and H1N1/H3N2 ferrets (Fig. 2b, c and Supplemental Fig. 2a, b). This was primarily due to an increase in CD8^+^ T cells, since CD4^+^ T-cell counts did not significantly differ from placebo animals. In BAL, mRNA/mRNA treatment enhanced both CD3^+^ and CD8^+^ T-cell counts compared to placebo ferrets. The effect of prime-boost with mRNA-Flu vaccination on T cell numbers in the BAL was less effective compared to a single influenza virus infection, as H1N1/mock-treated ferrets displayed higher CD3^+^ numbers compared to mRNA/mRNA ferrets. To determine if the increased T-cell counts correlated with increased IFNγ-responses, we performed a correlation analysis between population counts and IFNγ-ELISpot counts induced by H1N1 peptide pool stimulation. CD8^+^ T cell counts showed the strongest correlation with IFNγ-ELISpot responses, indicating that the IFNγ-response in the BAL and NT was mainly mediated by CD8^+^ T cells (Supplemental Fig. 3a, b).

**Figure 3:**
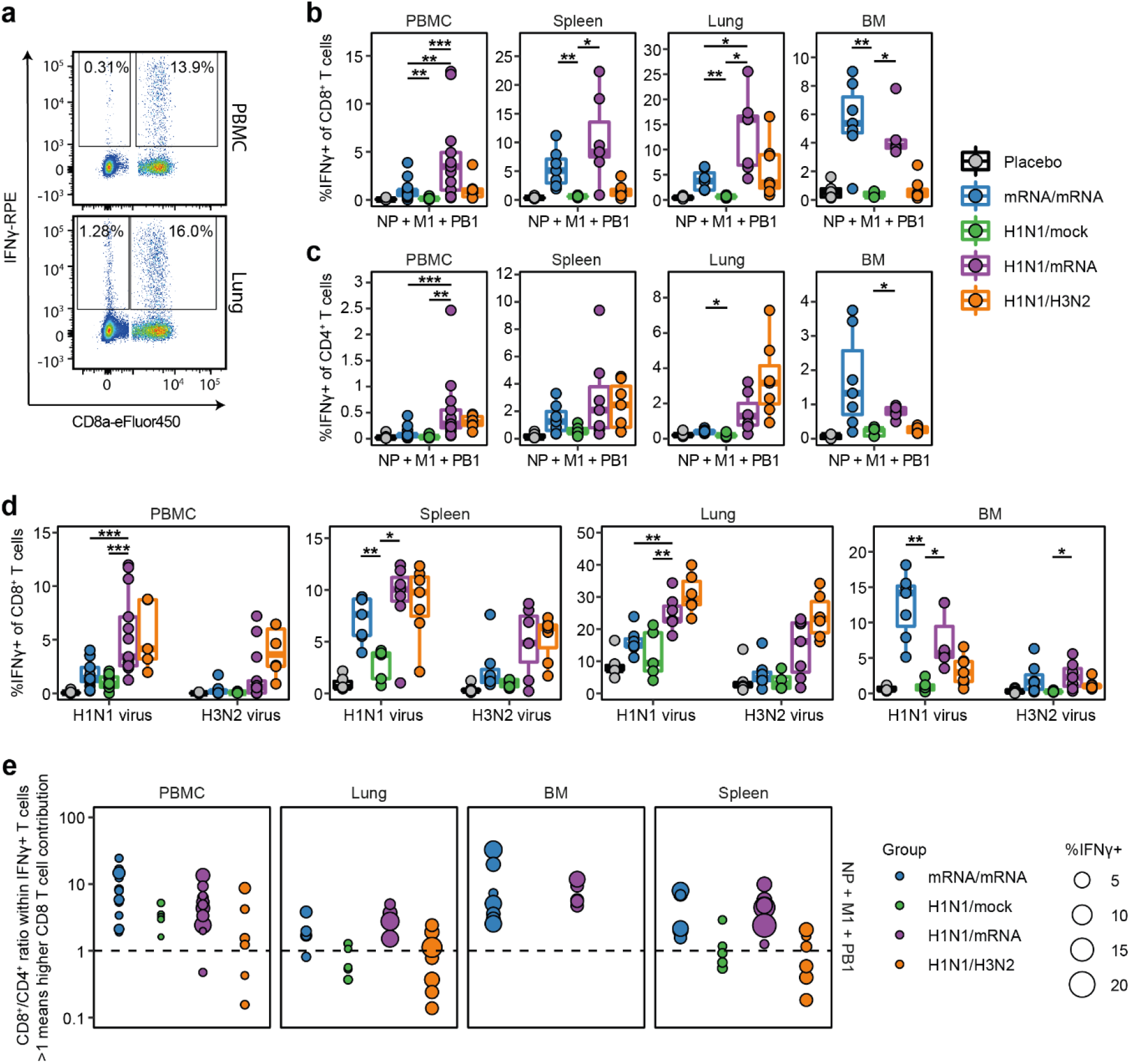
IFNγ responses of CD4^+^ and CD8^+^ T cells in PBMC, spleen, lung and bone marrow of immunized ferrets. Lymphocytes were stimulated with a peptide cocktail containing H1N1 NP, M1 and PB1 peptide pools or live influenza virus. Cells were stained for intracellular IFNγ and analyzed by flow cytometry. **a)** FACS plots depict representative CD4^+^ and CD8^+^ T-cell responses of H1N1/mRNA treated ferrets after peptide cocktail stimulation. Numbers indicate percentage of CD4^+^ or CD8^+^ T cells expressing IFNγ. **b, c)** IFNγ-positive CD8^+^ (B) and CD4^+^ (C) T cells after peptide cocktail stimulation. **d)** Percentage IFNγ-positive CD8^+^ T cells after stimulation with H1N1 (A/California/07/2009) or H3N2 (A/Uruguay/217/2007) influenza viruses. **e)** Ratio between CD8^+^ and CD4^+^ T cells within the CD3^+^ IFNγ^+^ T-cell population after peptide cocktail stimulation. Dotted line represents a ratio of 1 and samples with less than 50 CD3^+^ IFNγ^+^ cells were excluded from the analysis. Each dot represents one ferret and the dot size is relative to the total IFNγ response (%IFNγ^+^ of CD4^+^ and CD8^+^ T cells). Boxplots depict the median, 25% and 75% percentile, where the upper and lower whiskers extend to the smallest and largest value respectively within 1.5* the inter quartile ranges. In panels b-e, each dot represents one animal. n = 4-13 for PBMC and n = 4-7 for lung, spleen and BM. For visualization purposes, only comparisons between groups mRNA/mRNA, H1N1/mock and H1N1/mRNA are shown. No statistics were performed for panel e. An overview of all statistical comparisons is detailed in Supplemental data file 1. * = p < 0.05, ** = p < 0.01, *** = p < 0.001.

We additionally investigated cellular responses by IFNγ ELISpot in lungs that were perfused with a saline solution to reduce contamination of lung-derived lymphocytes with circulating lymphocytes. Remarkably, we observed robust cellular responses against NP, but not to M1 and PB1 in the lungs of mRNA/mRNA ferrets (Fig. 2d). Responses in the lung of mRNA/mRNA ferrets exceeded those measured in the blood, indicating that it is unlikely that the increase is due to contamination with circulating lymphocytes. In H1N1-primed ferrets, mRNA-Flu vaccination significantly boosted cellular responses against NP and M1 in the lung (group H1N1/mRNA vs H1N1/mock), to levels similar as achieved by a secondary natural infection with influenza virus (group H1N1/H3N2). Cellular responses against heterosubtypic virus stimulations (H3N2, H7N9, H5N1) were however similar between the H1N1/mRNA and mRNA/mRNA groups, indicating that mRNA/mRNA ferrets were not severely hampered by low responses against M1 and PB1 (Fig. 2d and Supplemental Fig. 1d).

Next, we investigated the presence of T-cell responses in the bone marrow (BM) since it is a reservoir for memory T cells [47]. mRNA/mRNA-treatment induced strong T-cell responses against NP in the BM (Fig. 2e and Supplemental Fig. 1d). Responses were similarly robust in H1N1/mRNA ferrets, while they were modest in H1N1/mock and H1N1/H3N2 ferrets. M1 and PB1 peptide pool responses were low for all groups in the BM, even though these responses were present in other tissues (Supplemental Fig. 4). The response to homologous (H1N1) and heterosubtypic (H3N2 [not significant for mRNA/mRNA], H5N1, H7N9) viruses was increased in both the mRNA/mRNA and H1N1/mRNA groups compared to H1N1/mock ferrets (Fig. 2e and Supplemental Fig. 1d). Together, these findings clearly demonstrate that the nucleoside-modified mRNA-LNP influenza T-cell vaccine is able to boost influenza virus-specific T-cell responses in the blood, respiratory tract and BM. Overall, compared to mRNA/mRNA ferrets, cellular responses were broader in H1N1/mRNA ferrets since they displayed robust M1 and PB1 responses in addition to NP (Supplemental Fig. 4).

**Figure 4:**
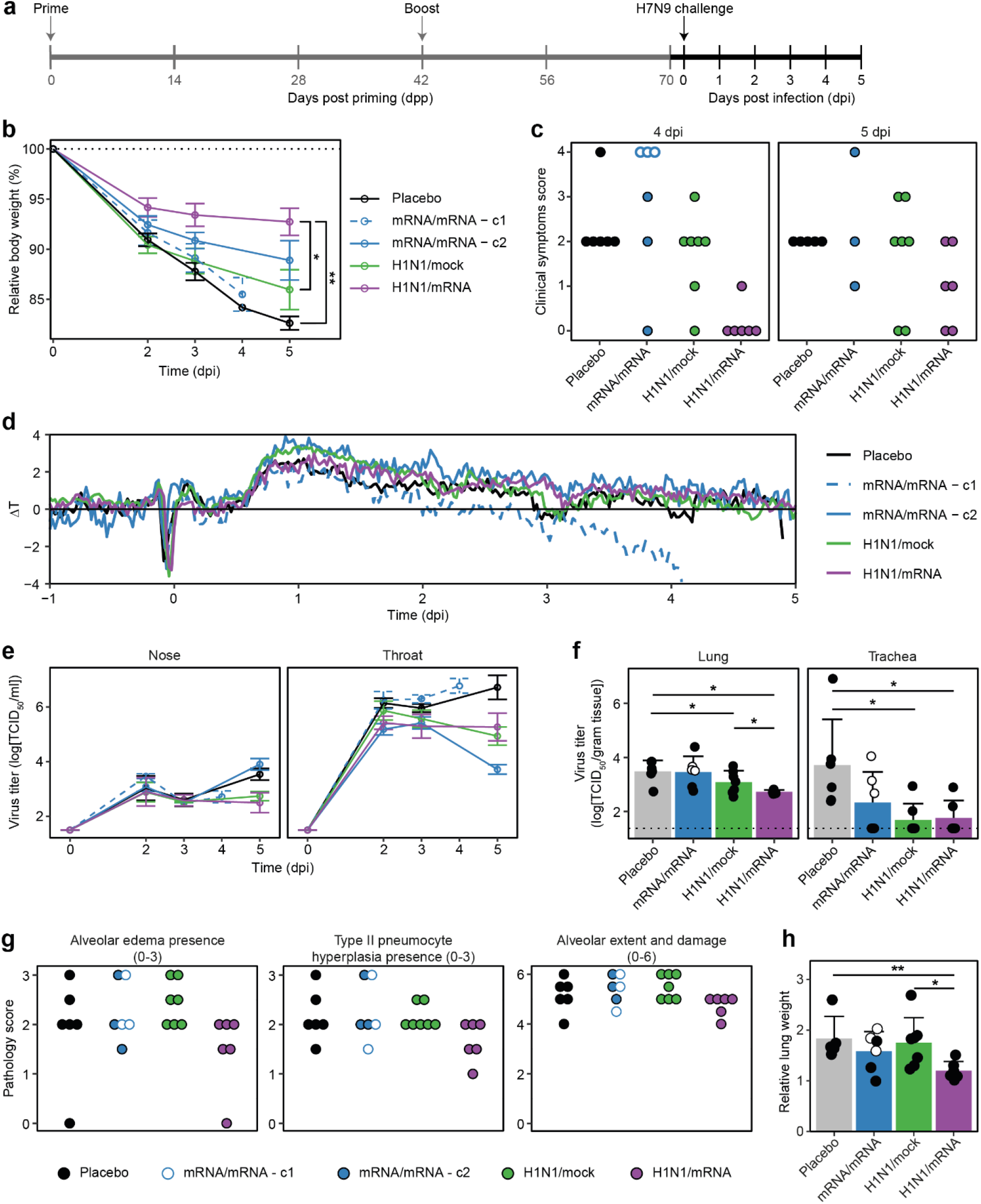
Boosting of existing immunity increases protection against H7N9 influenza virus challenge. **a)** Study layout depicting the H7N9 influenza virus challenge after different prime-boost regimens. Ferrets were challenged intratracheally with 10^6^ TCID_50_ A/Anhui/1/2013 (H7N9) influenza virus at 71 or 72 days post priming (dpp), which equals 0 days post infection (0 dpi). At 5 dpi, animals were euthanized after which pathology and virology was assessed. **b)** Decrease in body weight from 0 to 5 dpi. Body weight is depicted relative to body weight (%) on the day of challenge. **c)** Clinical scoring for parameters activity and breathing as detailed in the Materials & Methods. Ferrets reaching a combined score of 4 have reached the human endpoints and were euthanized. **d)** Fever depicted as temperature deviation from baseline. Baseline was determined as average body temperature from -5 to -1 dpi. **e, f)** Viral titers (TCID_50_) in **e)** nose and throat swabs and **f)** homogenized lung and trachea tissue as determined by endpoint titration on MDCK cells. Dotted line in panel f indicates the limit of detection. **g)** Pathology scoring for selected parameters as detailed in the Materials & Methods. **h)** Lung weight 5 dpi relative to body weight on the day of infection. For all panels n = 6-7. In panels b, e, f, and h, data are visualized as mean ± SD. In panel d, data is shown as group mean. In panels c and f-h, dots represent individual observations of ferrets. One placebo ferret and three mRNA/mRNA treated ferrets needed to be euthanized 4 dpi due to reaching the humane endpoints. The mRNA/mRNA ferrets euthanized 4 dpi are visualized as separate groups or depicted by open symbols (instead of filled). For visualization purposes, only comparisons between groups placebo, H1N1/mock and H1N1/mRNA are shown. No statistics were performed for panels c and g, as these are nominal data. An overview of all statistical comparisons is detailed in Supplemental data file 1. * = p < 0.05, ** = p < 0.01, *** = p < 0.001.

### The mRNA-based T-cell vaccine induces and boosts both CD4^+^ and CD8^+^ T-cell responses in PBMC, spleen, lung and bone marrow

To study the T-cell response in more detail, we measured IFNγ production of CD4^+^ and CD8^+^ T cells at 70 dpp by flow cytometric analysis. We stimulated lymphocytes derived from blood, spleen, lung and BM with an H1N1 peptide cocktail consisting of NP, M1 and PB1 peptide pools. mRNA/mRNA and H1N1/mRNA ferrets possessed significantly more CD8^+^IFNγ^+^ T cells in all tissues investigated relative to the placebo and H1N1/mock animals (Fig. 3a, b and Supplemental Fig. 5a, b). In PBMC and lung, H1N1/mRNA ferrets demonstrated significantly stronger CD8^+^ T-cell responses compared to mRNA/mRNA ferrets. Interestingly, the opposite was observed in the BM where mRNA/mRNA ferrets showed the most robust IFNγ-response, although this was not significantly stronger compared to H1N1/mRNA ferrets. Importantly, the H1N1 peptide cocktail-induced IFNγ-responses in PBMC, spleen and BM of H1N1/mRNA ferrets even exceeded those measured in ferrets boosted by a secondary infection (H1N1/H3N2 ferrets), further demonstrating the potency of the mRNA-Flu vaccine. In comparison to CD8^+^ T cells, CD4^+^ T-cell responses were weaker in most cases and differences between groups were slightly smaller (Fig. 3a, c and Supplemental Fig. 5a, c). Still, mRNA-Flu vaccination induced CD4^+^ T-cell responses in all investigated compartments of mRNA/mRNA ferrets and significantly boosted CD4^+^ T-cell responses in the blood and BM of H1N1/mRNA ferrets compared to H1N1/mock ferrets.

**Figure 5:**
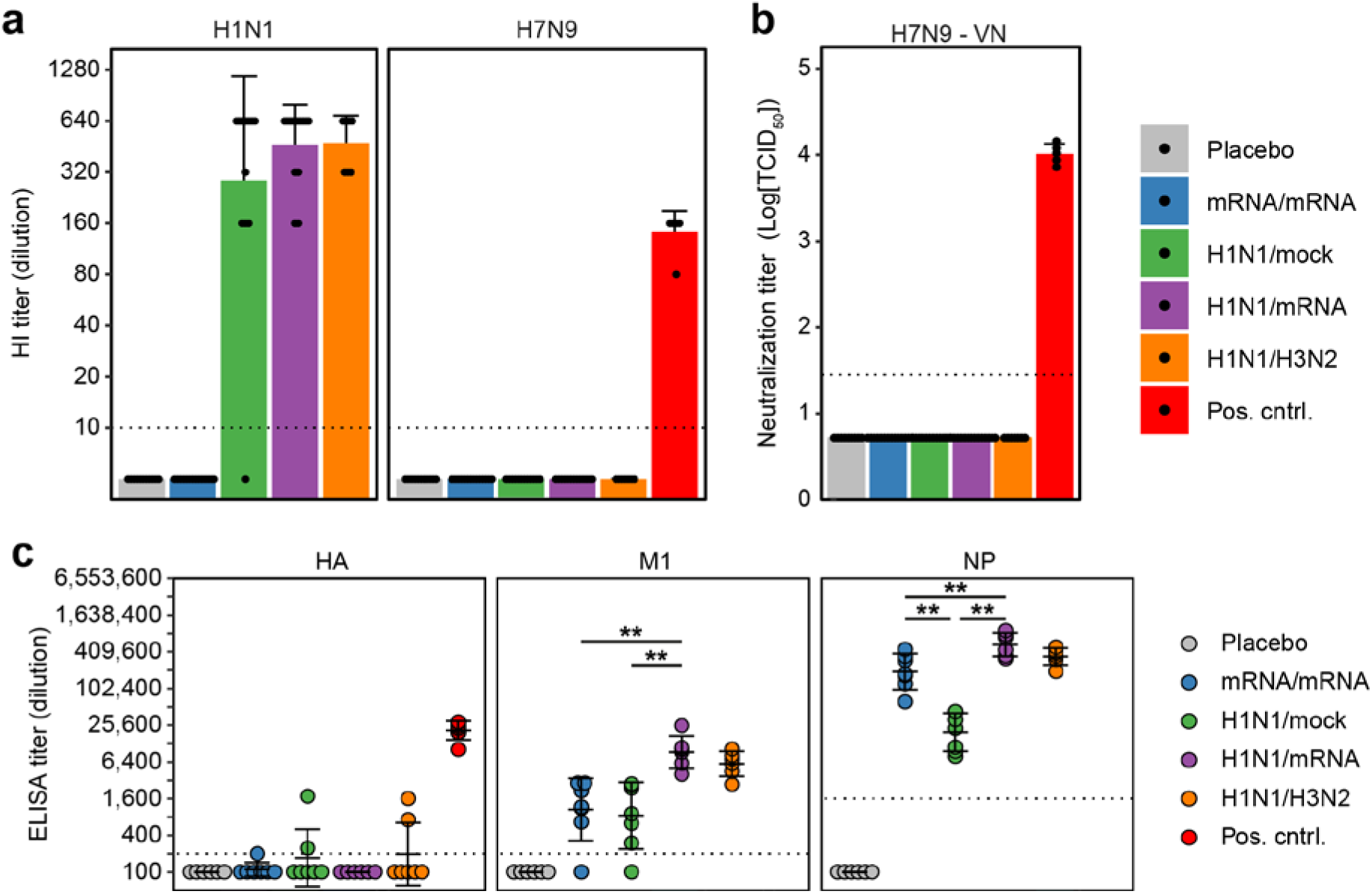
Antibody responses against H1N1 and H7N9 influenza viruses in sera obtained 70 days post priming (dpp). **a)** Antibodies against H1N1 (A/California/07/2009) or H7N9 (H7N9/PR8 reassortant) influenza virus, measured by hemagglutination inhibition (HI) assay. **b)** Virus neutralization titer against H7N9 influenza virus **c)** Antibodies binding to recombinant HA, M1 or NP of A/Anhui/1/2013 (H7N9) influenza virus measured by ELISA. The antibody titer is calculated as the extrapolated dilution of serum at which the OD 450 drops below background (mean of placebo animals + 3x SD). Positive control samples are sera from ferrets previously vaccinated twice with an H7N9 live attenuated virus [79]. Dotted line represents the lower limit of detection (a, b) or the background cut-off (c). In panels a-c, each dot represents one animal. For panels a and b, n = 7-14 (experimental groups) or n = 5 (positive control); for panel c, n = 6-7. For visualization purposes, only comparisons between groups mRNA/mRNA, H1N1/mock and H1N1/mRNA are shown in panel c. An overview of all statistical comparisons is detailed in Supplemental data file 1. * = p < 0.05, ** = p < 0.01, *** = p < 0.001.

Stimulations with live H1N1 or H3N2 influenza virus yielded similar results to H1N1 peptide cocktail stimulations. However, there was a trend that CD8^+^ T-cell responses in PBMC and lungs of H1N1/H3N2 ferrets were slightly stronger than in H1N1/mRNA ferrets (Fig. 3d). This is in part due to T cells that recognize conserved epitopes in proteins other than NP, M1 and PB1. CD4^+^ T-cell responses after virus stimulation were comparable to their CD8^+^ T cell counterparts, although CD4^+^ T-cell responses in the lung could not be interpreted because of high IFNγ background-responses in placebo animals (Supplemental Fig. 5d). Stimulations with H3N2 virus resulted in weaker CD4^+^ and CD8^+^ T cell responses compared to H1N1 virus stimulations (Fig. 3d and Supplemental Fig. 5d), which was not observed in the ELISpot assays (Fig. 2). This is likely due to a lower virus-to-cells ratio used for H3N2 stimulation in flow cytometry assays.

To investigate if mRNA-Flu vaccination leads to skewing of the T-cell response towards a CD4^+^ or CD8^+^ T-cell phenotype, we calculated the CD8^+^/CD4^+^-ratio within the CD3^+^IFNγ^+^ population after H1N1 peptide cocktail or H1N1 virus stimulation. In the tissues investigated, H1N1/mock and H1N1/H3N2 ferrets tended to have an average ratio of 1, demonstrating that IFNγ responses were approximately evenly distributed between CD4^+^ and CD8^+^ T cells (Fig. 3e and Supplemental Fig. 5e). Interestingly, in all tissues there was a clear skewing towards a CD8^+^ T-cell response in groups that received mRNA-Flu vaccination. Given the robust CD4^+^ T-cell responses in mRNA-Flu-immunized ferrets, skewing towards a CD8^+^ T cell response is not caused by a low CD4^+^ T-cell response, but by a very strong boosting of the CD8^+^ T-cell response. mRNA-Flu is thus a potent booster of both CD4^+^ and CD8^+^ T-cell immunity.

### H7N9 disease is reduced in influenza-experienced ferrets after booster vaccination

Next, we investigated whether mRNA-Flu vaccination could protect against severe disease caused by a heterosubtypic avian influenza virus infection. To this end, we immunized ferrets as described above with the exception of H1N1/H3N2 ferrets and challenged these animals intratracheally (i.t.) with a lethal dose of 10^6^ TCID_50_ H7N9 influenza virus four weeks after the booster vaccination (Fig. 4a). At this time, the boosted T-cell response is expected to be in its memory phase, similar to when (vaccinated) individuals are infected with influenza virus. Ferrets were euthanized five days post infection (dpi) to study viral replication and pathology.

Importantly, mRNA-Flu vaccination enhanced protection against H7N9 disease in H1N1-primed ferrets. Weight loss of H1N1/mRNA ferrets was limited to 7% and stabilized 5 dpi, while placebo animals lost more than 17% of bodyweight on average and were still losing weight at 5 dpi (Fig. 4b). mRNA/mRNA ferrets showed mixed results, with weight loss in isolator 1 being similar to placebo (∼15%) but less severe in isolator 2 (∼11%). Of note, one (out of six) placebo ferrets and three (out of six) mRNA/mRNA ferrets displayed inactivity and severe impaired breathing at 4 dpi and needed to be euthanized due to reaching the human end-points. The mRNA/mRNA group was clearly affected by a cage-effect of unknown origin as all ferrets that reached the humane endpoints were housed in one of the two isolators. The cage effect could not be explained by pre-existing immunity or infection history with other viruses (e.g. influenza virus, Aleutian disease, ferret corona viruses), as these were similar between groups (Supplemental Table 2). The two mRNA/mRNA groups are therefore analyzed together but visualized separately. No cage-effect was present in other treatment groups.

Weight data were in line with clinical symptoms as H1N1/mRNA-treated ferrets had less difficulty with breathing and were more active compared to other groups at 4 and 5 dpi (Fig. 4c). The height and duration of fever was not influenced by prior treatment as all groups displayed similar increases in body temperature (Fig. 4d, Supplemental Fig. 6a). Three animals in the mRNA/mRNA group showed hypothermia starting from 2 dpi and were euthanized at 4 dpi. Viral titers in nose and throat swabs were similar between groups at 2 and 3 dpi (Fig. 4e). By 5 dpi however, viral titers were lower in both H1N1/mRNA and H1N1/mock ferrets when compared to placebo. mRNA/mRNA ferrets gave mixed results. While viral titers in the nose were similar to placebo at all time-points investigated, viral titers in the throat at 5 dpi were significantly lower in surviving mRNA/mRNA ferrets compared to all other groups. We additionally measured viral titers in lung tissue. Differences were small, but H1N1/mRNA ferrets displayed significantly lower viral titers compared to all other groups (Fig. 4f). Viral titers in the trachea were low for all groups, except for the placebo group, indicating that all strategies limited viral replication to some extent.

Despite the reduced disease severity in H1N1/mRNA ferrets, the lungs showed moderate to severe broncho-interstitial pneumonia, often related to the bronchioles and bronchi which extended to the alveoli, irrespective of treatment (Fig. 4g, Supplemental Fig. 6b). However, alveolar edema, hyperplasia of Type II pneumocytes and alveolar damage was somewhat reduced in H1N1/mRNA-vaccinated ferrets. When we measured lung weight at 5 dpi as an independent measurement of lung pathology, H1N1/mRNA ferrets had significantly lower lung-weights (Fig. 4h). This indicates that inflammation and the resulting edema was less severe, which is in line with the less impaired breathing we observed in H1N1/mRNA ferrets. From these results, we conclude that nucleoside-modified mRNA-LNP influenza booster vaccination in H1N1-experienced ferrets was able to reduce H7N9 disease severity and virus replication.

### Protection against H7N9 influenza virus is likely mediated by cellular responses

To assess whether enhanced cellular responses during H7N9 influenza virus infection are related to the observed disease outcomes, we collected PBMCs at 4 or 5 dpi (depending on when ferrets were euthanized) and stimulated cells with H1N1 peptide pools in an IFNγ ELISpot assay. Although cellular responses against M1 and PB1 were low before infection (Figure 1b), they became more substantial after infection (Supplemental Fig. 7), suggesting that M1- and PB1-specific T cells may play a role in the observed reduction in H7N9 disease parameters. Differences between groups were difficult to quantify due to the strong responses observed, which reached the upper limit of detection of the IFNγ ELISpot assay.

To exclude the possibility that antibodies against H7N9 influenza virus played a role in the protection against H7N9 infection we measured the level of antibodies in ferret sera before H7N9 infection (70 dpp). We did not detect H7N9-specific antibodies by hemagglutination inhibition (HI) and virus neutralization (VN) assays (Fig. 5a, b). We additionally measured antibodies against H7N9 HA (H7), NP and M1 proteins by ELISA as not all influenza virus-specific antibodies can be detected by HI and VN assays. We did not find significant responses against H7, but we measured high antibody titers against NP and M1 (Fig. 5c). We could not investigate PB1-specific antibodies as the recombinant H7N9 PB1 protein was not commercially available. These findings indicate that HA-specific antibodies did not play a role in the disease reduction we observed, but the role of NP-, M1- and possibly PB1-specific antibodies remains to be investigated.

## Discussion

The COVID-19 pandemic has shown the enormous potential of the nucleoside-modified mRNA-LNP vaccine platform for inducing protective immune responses against SARS-CoV-2 infection in humans. This success is driving the development of mRNA-LNP vaccines against other infectious diseases, with influenza virus as a prime example. In fact, there are currently multiple mRNA-based influenza vaccines in the clinical phase of development [48]. Most of these vaccines are primarily focused on inducing neutralizing antibodies against the globular head domain of HA, which does not solve the problem of strain-specific immunity mediated by such antibodies. T cells could target a wider range of influenza viruses, but not much is known about the potential of mRNA-LNP vaccines to induce protective influenza-specific T-cell immunity. Here, we utilized a unique ferret model in which we could measure systemic and respiratory T-cell responses to evaluate the protective capacity of a nucleoside-modified mRNA-LNP vaccine encoding three conserved influenza proteins (mRNA-Flu). To our knowledge, this is the first study that provides a detailed evaluation of an mRNA-based influenza vaccine in a relevant animal model of influenza virus infection.

To mimic the human situation, we tested a combined nucleoside-modified mRNA-LNP vaccine (mRNA-Flu) encoding the internal influenza proteins NP, M1 and PB1 as a prime-boost strategy in naïve ferrets or as a booster in influenza-experienced ferrets. Prime-boost vaccination with mRNA-Flu resulted in robust, broadly-reactive cellular responses in blood, spleen, lung, NT and BM, although responses were primarily targeted against NP. mRNA-Flu was even more effective as a booster vaccination in influenza-experienced ferrets as it enhanced T-cell responses in all tissues investigated – including the BAL – and also boosted responses against M1 and PB1. To test the protective effect of the induced immune response, we challenged ferrets with avian H7N9 influenza virus as this strain has repeatedly transmitted from birds to humans and is considered as potentially pandemic [49]. After challenge, influenza-experienced ferrets that were boosted with mRNA-Flu lost less weight, showed fewer clinical symptoms and their lungs contained less edema compared to ferrets that did not receive an mRNA-Flu booster vaccination. We did not observe a similar protection for ferrets prime-boosted with mRNA-Flu only, which might be due to less robust and broad T-cell responses in the respiratory tract. Still, these results show that our nucleoside-modified mRNA-LNP T-cell vaccine is a promising candidate to boost broadly-reactive cellular responses and can be used to enhance protection against heterosubtypic influenza viruses.

To induce a broadly-reactive T-cell response, we developed a vaccine targeting three immunogenic conserved influenza proteins. We have previously shown that both ferrets and healthy human blood donors possess clearly detectable NP-, M1- and PB1-reactive T-cells [25]. In our current experimental model, both a single mRNA-Flu vaccination and H1N1 influenza virus infection elicited NP-specific responses. Responses against M1 and PB1 were weaker, especially in mRNA-Flu-primed animals.

However, booster vaccination increased M1- and PB1-specific responses in all H1N1-primed ferrets and approximately half of the mRNA/mRNA ferrets developed detectable M1 and PB1-specific responses. Although it is unclear why M1- and PB1-responses were weak initially, these responses substantially increased shortly after H7N9 influenza virus challenge, suggesting that M1- and PB1-specific T cells played a role in reducing H7N9 influenza disease. This indicates that it could be beneficial if future mRNA-based influenza vaccines targeted multiple relatively well-conserved internal proteins. This would also safeguard against influenza virus mutations as the virus is less likely to escape from a broad immune response.

The T cells induced by mRNA-Flu vaccination responded to a wide range of influenza viruses, including seasonal H3N2, pandemic H2N2 and avian H5N1 and H7N9 strains. Previous research has already shown that T cells are crucial for protection against heterosubtypic infections, especially lung resident memory T-cells (Trm) [46, 50]. We show that mRNA-LNP vaccination – in contrast to inactivated influenza vaccines [51] – is able to induce T cells residing in the respiratory tract, even when given i.m. Whether these T cells possess a Trm phenotype still remains to be elucidated due to a lack of ferret-specific reagents. The T-cell responses we found in NT and lung after mRNA-Flu prime-boost confirms a previous report of Lackzo et al. who found that i.m. administration of mRNA-LNP vaccines induced potent cellular responses in the lungs of mice [52]. The responses we found were not an artefact of circulating lymphocytes as lungs were perfused and cellular responses in the lung were higher than those in the blood, showing that influenza virus-specific T cells accumulated in the lung tissue. Still, responses in the BAL were absent in mRNA-Flu prime-boosted ferrets, indicating that local presentation of antigen and/or inflammation is required for extended tissue-residing cellular immunity. Intranasal administration of mRNA vaccines could potentially enhance protection by also inducing T cells in the BAL and increasing T-cell numbers in the NT, but additional research needs to be performed to overcome the epithelial barrier and to prevent excessive immune activation [53]. Remarkably, mRNA-Flu vaccination boosted cellular responses in the BAL, NT and lungs of H1N1-primed ferrets that reacted not only to NP, but also to M1 and PB1. This is a particularly relevant finding as a large part of the human population has already been naturally exposed to influenza virus. For this group, a single mRNA-LNP immunization administered i.m. might be sufficient to boost respiratory T-cell responses. These findings stress the importance of animal models that reflect the human infection history as pre-existing immunity can clearly influence vaccine responses.

mRNA-Flu also induced potent responses in the BM. This might be partly caused by the close proximity of mRNA-Flu administration (hind legs) and T-cell isolation from the BM (femur). In fact, T cells can be primed in the BM after local antigen presentation [54, 55]. This can be beneficial for the longevity of the cellular response as the BM serves as a reservoir for memory T cells [56, 57]. The observation that nucleoside-modified mRNA-LNP vaccination is a potent inducer of BM-residing T-cell immunity warrants further investigations into the longevity and importance of this response.

In our study, vaccine-induced T-cell responses consisted of both CD4^+^ and CD8^+^ T cells. Similarly, Freyn et al. found that a single dose of H1N1 NA- or NP-encoding mRNA-LNP induced robust CD4^+^ and CD8^+^ T-cell responses in mice [40]. In humans, SARS-CoV-2 mRNA-LNP vaccines also induced both CD4^+^ and CD8^+^ T cells, although the extent to which the vaccines induced CD4^+^ and CD8^+^ T cells differs between studies [34, 58, 59]. We found that the T-cell response after mRNA-Flu booster vaccination was skewed towards a CD8^+^ phenotype. This skewing might be beneficial, as clearing off virus-infected cells is primarily mediated by CD8^+^ T cells [20]. It should be mentioned, however, that we could only measure IFNγ responses and we might have missed activated CD4^+^ T cells that responded by producing other typical CD4^+^ cytokines such as TNF-*α* and IL-2.

Besides T cells, the mRNA-Flu vaccine also induced humoral responses against NP, M1 and possibly PB1; antibodies against PB1 could not be measured due to the lack of reagents. We did not find any functional role for NP- and M1-antibodies by HI and VN assays, although these assays primarily detect (neutralizing) anti-HA antibodies. Still, in mice, vaccination with recombinant NP induced potent anti-NP antibodies that protected against severe disease after an influenza virus challenge, but only if these mice also possessed functional T cells [60, 61]. This protection might be mediated by antibody-dependent cell cytotoxicity (ADCC) activity, although it is still uncertain if NP- and M1-specific antibodies can facilitate ADCC [62, 63]. Whether ADCC or other effector mechanisms played a role in our study remains therefore unknown. Future serum transfer experiments in ferrets could help in clarifying the exact role of NP-, M1- and PB1-specific antibodies in the protection against influenza virus disease.

To evaluate the robustness of T-cell-mediated protective immunity, we utilized a ferret challenge model in which a lethal dose of H7N9 influenza virus was deposited directly into the lungs of animals by intratracheal inoculation. In this way, a large amount of pneumocytes become directly infected and T cells are only granted a short timeframe to become activated and prevent further disease. This robust challenge model is not representative of a normal human exposure. People typically encounter a lower viral load [64] and primarily in the upper respiratory tract, which affords T cells a longer time to establish protective immunity. We thus expect a greater protective effect of the T-cell response upon natural infection doses. The challenge model we used – while not utilizing a natural inoculation route and dose – very well represents the severe pneumonia observed in humans hospitalized with H7N9 influenza virus infection, which cannot be achieved with lower infection doses and other inoculation routes.

We could not clearly establish whether a prime-boost strategy with mRNA-Flu was protective likely due to a cage effect. Ferrets prime-boosted with mRNA-Flu housed in one isolator showed protection against H7N9 influenza disease similarly to mRNA-Flu-boosted influenza-experienced ferrets. Ferrets in the second isolator however showed more severe symptoms after infection than the placebo animals and needed to be euthanized one day prior to the scheduled termination of the experiment. We did not find differences between the two cages that explain this discrepancy. Both humoral and cellular immune responses were similar, ferrets tested negative for Aleutian disease and showed similar previous exposure to canine distemper virus and ferret corona viruses. For practical reasons, the H7N9 influenza virus challenge was performed on two consecutive days with each treatment group split over both days (see Materials & Methods for details). It is unlikely that differences are due to separate preparation of the inoculum, as all other groups – which were also divided over two days – did not respond differently to the challenge. Additional experiments would be required to clarify if the influenza-specific T-cell response induced by prime-boost vaccination with mRNA-Flu is protective in naïve ferrets.

In contrast to traditional inactivated influenza virus vaccines, nucleoside-modified mRNA-LNP vaccines can induce both humoral and cellular immunity [34-38]. With the induction of a broadly-reactive T-cell response, these vaccines should be less sensitive to antigenic drift and shift that have hampered traditional HA-based vaccines. Furthermore, mRNA-LNP SARS-CoV-2 vaccines perform remarkably well in elderly people [65, 66], while inactivated influenza virus vaccines often have subpar performance with increasing age [67]. mRNA-based influenza vaccines might thus be especially suited to protect this group that is at high risk for influenza-related mortality and morbidity. For these reasons, the nucleoside-modified mRNA-LNP platform is a viable option for the improvement of seasonal influenza vaccination. The inclusion of conserved internal influenza virus proteins could additionally provide protection against potential pandemic influenza viruses, as demonstrated in the current study. To our best knowledge, this is the first study that provides a detailed evaluation of an mRNA-based combined influenza T-cell vaccine in a highly relevant ferret model. We postulate that the nucleoside-modified mRNA-LNP-based influenza vaccine can boost the number of broadly-reactive T-cells to a level that prevents severe disease and death, reducing the impact of future influenza epidemics and pandemics on the society.

## Acknowledgements

We would like to thank Marion Hendriks, Jolanda Kool, Helena Pinheiro Guimarães, Noortje Smits, Ronald Jacobi and Martijn Vos for their help during the animal sections, the biotechnicians from the animal facility for excellent care-taking of the animals and Dr. Teun Guichelaar and Dr. Willem Luytjes for critical reviewing of the manuscript.

## Author contributions

Conceptualization: KvdV, DvB, NP, JdJ

Methodology: KvdV, JdJ

Software: JAF

Formal analysis: KvdV, JvdB

Investigation: KvdV, JL, HvD, CVBdM, SL, FP

Resources: NP, HM, MBB, PJCL

Data curation: KvdV

Writing – original draft: KvdV, JAF

Writing – review & editing: JL, DvB, NP, JdJ

Visualization: KvdV

Supervision: JdJ

Project administration: JdJ

Funding acquisition: JdJ, NP

## Materials & Methods

### Ethics statement

The experiment was approved by the Animal Welfare Body of Poonawalla Science Park – Animal Research Center (Bilthoven, The Netherlands) under permit number AVD3260020184765 of the Dutch Central Committee for Animal experiments. All procedures were conducted according to EU legislation. Ferrets were examined for general health on a daily basis. If animals showed severe disease according to the defined end points prior to scheduled termination they would be euthanized by cardiac bleeding under anesthesia with ketamine (5 mg/kg; Alfasan, Woerden, The Netherlands) and medetomidine (0.1 mg/kg; Orion Pharma, Espoo, Finland). Endpoints were scored based on clinical parameters for activity (0 = active; 1 = active when stimulated; 2 = inactive and 3 = lethargic) and impaired breathing (0 = normal; 1 = fast breathing; 2 = heavy/stomach breathing). Animals were euthanized when they reached score 3 on activity level (lethargic), when the combined score of activity and breathing impairment reached 4 or if their body weight decreased by more than 20%.

### Cell & virus culture

MDCK cells were grown in MEM (Gibco, Thermo Fisher Scientific, Waltham, MA) supplemented with 10% fetal bovine serum (FBS; HyClone, GE Healthcare, Chicago, IL), 40 μg/ml gentamicin and 0.01M Tricin (both from Sigma-Aldrich, Saint Louis, MO). VERO E6 cells were cultured in DMEM (Gibco) supplemented with 10% FBS and 1x penicillin-streptomycin-glutamine (Gibco). A/California/07/2009 (H1N1), A/Switzerland/97-15293/2013 (H3N2), A/Vietnam/1203/2004 WT (H5N1), A/Anhui/1/2013 (H7N9) and H7N9/PR8 reassortant (NIBRG-268, NIBSC code 13/250) influenza viruses were obtained from the National Institute for Biological Standards and Control (NIBSC, Hertfordshire, England). Influenza virus was grown on MDCK cells in MEM medium supplemented with 40 µg/ml gentamicin, 0.01M Tricine and 2 µg/ml TPCK treated trypsin (Sigma-Aldrich). At >90% cytopathic effect (CPE), the suspension was collected and spun down (4000x g for 10 minutes) to remove cell debris. H1N1 and H3N2 virus was sucrose purified on a discontinuous 10-50% sucrose gradient. Due to BSL-3 classification of H7N9 and H5N1, the virus was not purified. All virus aliquots were snap-frozen and stored at -80 °C.

### mRNA production

NP, M1 and PB1 mRNAs are based on the A/Michigan/45/2015 H1N1pdm virus, which is nearly identical to A/California/07/2009 (NP = 99.2%, M1 = 98.4% and PB1 = 99.6% conserved). Production of mRNAs was performed as described earlier [40, 68]. Briefly, codon-optimized NP, M1, and PB1 genes were synthesized (Genscript, Piscataway, NJ) and cloned into an mRNA production plasmid. T7-driven in vitro transcription reactions (Megascript, Ambion, Thermo Fisher) using linearized plasmid templates were performed to generate mRNAs with 101 nucleotide long poly(A) tails. Capping of mRNAs was performed in concert with transcription through addition of a trinucleotide cap1 analog, CleanCap (TriLink, San Diego, CA) and m1Ψ-5’-triphosphate (TriLink) was incorporated into the reaction instead of UTP. Cellulose-based purification of mRNAs was performed as described [69]. mRNAs were then tested on an agarose gel before storing at -20 °C.

### Lipid nanoparticle formulation of mRNA

Purified mRNAs were formulated into lipid nanoparticle using a self-assembly process wherein an ethanolic lipid mixture of an ionizable cationic lipid, phosphatidylcholine, cholesterol, and polyethylene glycol-lipid was rapidly combined with an aqueous solution containing mRNA at acidic pH as previously described [45]. The ionizable cationic lipid (pKa in the range of 6.0-6.5, proprietary to Acuitas Therapeutics, Vancouver, Canada) and LNP composition are described in the patent application WO 2017/004143. The average hydrodynamic diameter was ∼80 nm with a polydispersity index of 0.02-0.06 as measured by dynamic light scattering using a Zetasizer Nano ZS (Malvern Instruments Ltd, Malvern, UK) and an encapsulation efficiency of ∼95% as determined using a Ribogreen assay.

### Animal handling

63 female ferrets (*Mustela putorius furo*) aged 12-13 months (Euroferret, Copenhagen, Denmark) were delivered three weeks before commencement of the study and were semi-randomly distributed by weight. Ferret throat swabs were screened for SARS-CoV-2 by RT-qPCR as described before [70] and ferret sera was screened for influenza exposure by NP ELISA (Innovate Diagnostics, Grabels, France) and HI. Additionally, ferret sera (ELISA) and swabs (RT-qPCR) were screened for other corona viruses, canine distemper virus and Aleutian disease by the European Veterinary Laboratory (EVL, Woerden, the Netherlands). All ferrets tested negative for influenza and SARS-CoV-2; four animals displayed low antibody titers against Aleutian disease; all animals possessed titers for CDV-antibodies but tested negative for active infection by RT-qPCR. Ferrets were housed per 3 or 4 animals in open cages and received pelleted food (Altromin 5539) and water *ad libitum*. Animals were visually inspected daily and weighed at least once per 7 days. Light was adjusted to 9.5 hours per day to prevent the ferrets from going into estrous. For influenza infections animals were moved to BSL-3 level isolators. Due to a limited number of isolators, groups that did not receive an infection were kept housed in regular open cages. 14 days after infection the animals were confirmed to be negative for infectious influenza and moved back to regular housing.

Ferrets that received a (mock) infection were swabbed and weighed at 0, 2, 4, 7, 9 and 14 days after the first and second infection. Vaccinated animals were only swabbed at days 0 and 14 and weighed on days 0, 7 and 14. Blood was collected from the vena cava at 0, 14, 28, 42, 56 and 71 days post priming (dpp). These handlings were performed under anesthesia with ketamine (5 mg/kg). Blood was collected by heart puncture on 70 and 76 dpp. Infections, vaccinations, temperature transponder implantation and euthanasia were performed after anaesthetization with ketamine and medetomidine (0.1 mg/kg). Animals that received a temperature transponder (Star Oddi, Garðabær, Iceland) abdominally received 0.2 ml Buprenodale (AST Farma, Oudewater, The Netherlands) as a post-operative analgesic. Anesthesia with medetomidine was antagonized with atipamezole (0.25 mg/kg; Orion Pharma), but was delayed by 30 minutes in case of infection/vaccination to prevent sneezing and coughing.

### Study outline

The study consisted of five experimental groups: 1) placebo; 2) mRNA/mRNA; 3) H1N1/mock; 4) H1N1/mRNA; and 5) H1N1/H3N2. Each experimental group consisted of 14 (group 1-4) or 7 (group 5) ferrets. For practical reasons the experiment was split into three sub experiments (A, B and CD). All sub-experiments followed the same regime up to day 70 of the experiment, but were started 8 days after each other. Sub-experiments A and B both contained groups 1-5 with 3-4 animals/group and were terminated 70 dpp to study the immune response. Sub-experiment CD contained groups 1-4 with 7 animals/group, split over 2 cages. Sub-experiment CD was again dived into two smaller sub-experiments (C and D) on 71 dpp, which were challenged with H7N9 on 71 and 72 dpp respectively. Data from the different sub-experiments were visualized and analyzed together.

On day 0, groups 3-5 were inoculated intranasally (i.n.) with 10^6^ TCID_50_ H1N1 in 0.1 ml inoculum. Group 1 received PBS in the same manner. Group 2 was administered 250 μl of mRNA vaccine – containing 50 μg of NP, M1 and PB1 – in their left or right hindleg. On 42 dpp, animals received a booster treatment. Group 5 was inoculated i.n. with 10^6^ TCID_50_ H3N2 in 0.1 ml inoculum. Group 1 was treated similarly but received PBS instead of H3N2 virus. Groups 2-4 were injected with 250 μl of influenza-mRNA vaccine (groups 2, 4) or Luciferase-mRNA (group 3; 50 μg) in their left or right hindleg. At 70 dpp, seven ferrets of each group were euthanized to study the immune response in the respiratory tract. The other seven animals (excluding group 5) were challenged intratracheally (i.t.) with 10^6^ TCID_50_ H7N9 in 3ml inoculum at 71 or 72 dpp. Five days later, ferrets were euthanized to study viral titers and pathology.

Animals were euthanized by heart puncture and blood and serum was collected. For ferrets in sub-experiments A and B, the lungs were perfused as described before [25] and broncho-alveolar lavage (BAL) was collected by flushing the lungs twice with 30ml of room temperature (RT) RMPI1640 (Gibco). The BAL fluid was then kept on ice till processing. Lungs, spleen, femur (right leg) and nasal turbinates (NT) were collected in cold RMPI1640 supplemented with 10% FBS and 1x penicillin-streptomycin-glutamine and stored at +4 °C until processing. For ferrets in sub-experiments C and D, lungs were weighed before the left cranial and caudal lobes were inflated with and stored in 10% formaldehyde for later pathological analysis. Small slices of the right cranial, middle and caudal lobes were put in Lysing matrix A tubes (MP Biomedicals, Irvine, CA) and stored at -80 °C until later virological analysis. The lower part of the trachea was stored in 10% formaldehyde for pathology and 1 cm of the middle part of the trachea was stored in Lysing matrix A tubes.

### Tissue processing

Blood was collected in 3.5 ml VACUETTE tubes with clot activator (Greiner, Merck, Kenilworth, NJ) and spun down at 4000x g for 10 minutes to isolate serum. Heparin blood was collected in 9 ml sodium-heparin coated VACUETTE tubes (Greiner) and diluted 1:1 with PBS (Gibco) for density centrifugation on a 1:1 mixture of LymphoPrep (1.077 g/ml, Stemcell, Vancouver, Canada) and Lympholyte-M (1.0875 g/ml, Cedarlane, Burlington, Canada). Cells were spun for 30 minutes at 800x g, after which the interphase was collected and washed thrice with washing medium (RPMI1640 + 1% FCS + 1x penicillin-streptomycin-glutamine). Next, cells were resuspended in stimulation medium (RPMI1640 + 10% FCS + 1x penicillin-streptomycin-glutamine) and counted using a hemocytometer.

Spleen, lung and NT tissue were processed as detailed before [25]. In brief, spleens were homogenized in a sieve using the plunger of a 10 ml syringe. The resulting suspension was collected while excluding the larger debris and pelleted by centrifugation for 10 minutes at 500x g. The pellet was resuspended in 50 ml EDTA-supplemented (2mM) washing medium and transferred over a 100 µm SmartStrainer (Miltenyi Biotec, Bergisch Gladbach, Germany). The cell suspension was then diluted to 90 ml, which was divided into 3x 30 ml and layered on top of 15 ml Lympholyte-M for density centrifugation similar to that of blood. All washing steps were performed with EDTA-supplemented medium to prevent agglutination of cells.

Lungs were cut into 5 mm^3^ cubes and digested in 12 ml of collagenase I (2.4 mg/ml, Merck) and DNase I (1 mg/ml, Novus Biologicals, Centennial, CO) for 60 minutes at 37 °C while rotating. Samples were homogenized in a sieve using a plunger, spun down for 10 minutes at 500x g and resuspended in washing medium. This suspension was transferred over a 70 μm cell strainer (Greiner) and used for density centrifugation similar to that of the spleen.

Nasal turbinates were mashed on a sieve using a plunger and pelleted by spinning for 5 minutes at 500x g. The pellet was resuspended in 3 ml collagenase/DNAse solution (similar to lung) and incubated for 30 minutes at 37 °C while rotating. Next, the suspension was directly mashed over a 70 μm cell strainer (Greiner) with a plunger and washed twice with 10 ml washing medium. The resulting pellet was resuspended in 6 ml of 40% Percoll (GE Healthcare) and layered on top of 70% Percoll to isolate leukocytes. Samples were spun for 20 minutes at 500x g after which the interphase was collected and washed twice with washing medium. After the final wash, cells were resuspended in stimulation medium and used for ELISpot and FACS.

After collection, 3 ml BAL was used for ELISpot without further processing. The remaining volume was spun down at 500x g for 5 minutes and resuspended in 12 ml FACS buffer (PBS [Gibco]+ 0.5% BSA [Merck] + 2mM EDTA). The suspension was transferred over a 70 μm SmartStrainer (Miltenyi Biotec), spun down at 500x g for 5 minutes and resuspended in FACS buffer. This suspension was used for FACS.

Femurs were cleaned from residual tissues and briefly decontaminated with 70% ethanol. The femur was then cut on both sides so that the shaft could be flushed with 15 ml of ice-cold RPMI washing medium. The suspension was transferred over a 70 μm cell strainer and pelleted by centrifugation for 7 minutes at 500xg at 4 °C. Erythrocytes were lysed with ACK lysis buffer after which the suspension was spun down, resuspended in washing medium and again transferred over a 70 μm cell strainer. The resulting suspension was spun down, resuspended in stimulation medium and used for ELISpot and FACS.

### Peptide pools

NP (NR-18976), M1 (NR-21541) and PB1 (NR-18981) H1N1 peptide arrays were obtained through BEI Resources, NIAID, NIH. Peptides were supplied as individual aliquots and were pooled in-house after dissolving in H_2_O, 50% acetonitrile or DMSO depending on the solvability. The merged peptide-suspension was then aliquoted and speed-vacced for 48 hours to reduce the volume. Vials were stored at -80 °C.

H2N2 peptide pools were based on A/Leningrad/134/17/1957 and were custom ordered from JPT Peptide Technologies GmbH (Berlin, Germany). Each pool contained 15 amino acid long peptides with an overlap of 11 amino acids spanning the entire protein of NP, M1 or PB1. Peptides were synthesized as reported before [25]. HIV-1 Con B gag motif peptide pool (JPT) served as a negative control for our assays and was handled in the same way as the H2N2 peptide pools.

Before use, H1N1 and H2N2 peptide pools were dissolved in DMSO, aliquoted and stored at -20 °C. On the day of use, peptide pool aliquots were thawed and diluted with stimulation medium. The peptide pool suspension was added to cells, such that a final peptide concentration of 1 µg/ml per peptide with a DMSO concentration of less than 0.2% was achieved.

### ELISpot

Pre-coated Ferret IFNγ ELISpot (ALP) plates (Mabtech, Nacka Strand, Sweden) were used according to the manufacturers protocol. Lymphocytes were stimulated with live virus (MOI 100 for H3N2; MOI 1 for H5N1; MOI 0.1 for H1N1 and H7N9) or peptide pools (1 µg/ml) in ELISpot plates at 37 °C. Per well, 250K cells (PBMC), 400K cells (BM), 62.5K cells (lung lymphocytes) or undiluted cell suspension (BAL, nasal turbinates) was added. On day 56 – 2 weeks after booster vaccination – 125K PBMCs were used for viral stimulations due to high cellular responses. After 20 hours the plates were developed according to the manufacturers protocol, with the modification that the first antibody staining was performed overnight at 4 °C. Plates were left to dry for 2-3 days after which they were packaged under BSL-3 conditions and heated to 65 °C for 3 hours to inactivate any remaining infectious influenza particles. Analysis of ELISpot plates was performed using the ImmunoSpot® S6 CORE (CTL, Cleveland, OH).

### Flow cytometry – cell counts

BAL and NT samples were stained in 96-wells plates using the FoxP3 / Transcription factor staining buffer set (eBioscience, Thermo Fisher). Cells were stained with α-CD4-APC (02, Sino biological, Beijing, China), α-CD8a-eFluor450 (OKT8, eBioscience), α-CD14-PE (Tük4; Thermo Fisher) and Fixable Viability Stain 780 (BD, Franklin Lakes, NJ) in 100 μl for 30 minutes at 4^°^C. Samples were then washed twice with 150 μl FACS buffer, followed by fixation with 100 μl fixative from the FoxP3 staining kit for 20 minutes at RT. Next, samples were washed twice with 150 μl 1x permeabilization buffer (FoxP3 staining kit). After the second wash, samples were stained with 100 μl permeabilization buffer containing α-CD3e-FITC (CD3-12, Bio-Rad, Hercules, CA) for 30 minutes at 4 °C. Samples were then washed twice with 150 μl 1x permeabilization buffer and once with 150 μl FACS buffer. After the last wash, samples were resuspended in 180 μl FACS buffer after which 50 μl precision count beads (Biolegend, San Diego, CA) were added to BAL and NT samples. Samples were measured in plates using the high-throughput system of a Symphony A3 system (BD). Data was analyzed using FlowJo™ Software V10.6 (BD).

### Flow cytometry – intracellular cytokine staining

Lymphocytes derived from blood, lung or BM were stimulated in U-bottom plates with 1-3 million cells/well. Stimulations consisted of medium, H1N1 live virus (MOI 1), H3N2 live virus (MOI 10), an H1N1 peptide cocktail containing peptide pools of NP, M1 and PB1 (1 μg/peptide/ml), and a HIV peptide pool (1 μg/peptide/ml) serving as a negative control. Cells were stimulated for 20 (virus, medium) or 6 hours (peptide pools) at 37 °C. During the last 5 hours of stimulation, 1x brefeldin A (Biolegend) was added to each well. Plates were then stored at 4 °C until they were stained the following morning. Staining and acquisition followed the same procedure as detailed above, with the exception that α-CD14-PE was absent in the extracellular staining and instead, α-IFNγ-RPE (CC302, MyBioSource, San Diego, CA) was added to the intracellular staining.

### TCID_50_ determination

Nose and throat swabs were collected in 2 ml transport medium containing 15% sucrose (Merck), 2.5 µg/ml Amphotericin B, 100 U/ml penicillin, 100 µg/ml streptomycin and 250 µg/ml gentamicin (all from Sigma) and stored at -80 °C. For analysis, swabs were thawed, vortexed, serially diluted and tested in sextuplicate on MDCK cells. Trachea and lung samples stored in Matrix A tubes were thawed and 750 μl of DMEM infection medium (DMEM containing 2% FBS and 1x penicillin-streptomycin-glutamine) was added. Tissues were then dissociated in a FastPrep-24™ by shaking twice for 1 minute after which the samples were spun down for 5 minutes at 4000x g. To determine viral titers, the supernatant was serially diluted in sextuplicate on MDCK cells. Cytopathic effect (CPE) was scored after 6 days of culturing and TCID_50_ values were calculated using the Reed & Muench method. Viral titers in virus stocks were similarly tested, but in octuplicate.

### ELISA

Immulon 2 HB 96-well plates (Thermo Fisher) were coated overnight at RT with 100 μl/well recombinant HA (0.5 μg/ml), NP (0.5 μg/ml) or M1 (0.25 μg/ml) protein of A/Anhui/1/2013 (Sino biologicals). The next day, plates were washed thrice with PBS + 0.1% Tween-80 before use. Sera were diluted 1:100 in PBS + 0.1% Tween-80 and then 2-fold serially diluted. Per well, 100 μl of diluted sera was added and plates were incubated for 60 minutes at 37 °C. After washing thrice with 0.1% Tween-80, plates were incubated for 60 minutes at 37 °C with HRP-conjugated goat anti-ferret IgG (Alpha Diagnostic), diluted 1:5000 in PBS containing 0.1% Tween-80 and 0.5% Protivar (Nutricia, Hoofddorp, The Netherlands). Plates were then washed trice with PBS + 0.1% Tween-80 and once with PBS, followed by development with 100 μl SureBlue™ TMB (KPL, Gaithersburg, MD) substrate. Development was stopped after 10 minutes by addition of 100 μl 2M H_2_SO_4_ and OD_450_-values were determined on the EL808 absorbance reader (Bio-Tek Instruments). Individual curves were visualized using local polynomial regression fitting with R software v4.1.1 [71]. Antibody titers were determined as the dilution at which antibody responses dropped below background. This background was calculated as the ‘mean + 3 * standard deviation’ of the OD_450_ at a 200x (HA, M1) or 1600x (NP) serum-dilution of placebo animals.

### Hemagglutination inhibition assay

Hemagglutination inhibition (HI) titers in ferret sera were determined in duplicate according to WHO guidelines [72]. In brief, sera were heat-inactivated at 56 °C for 30 minutes and treated with receptor destroying enzyme (Sigma) in a 1:4 mixture (5x dilution of sera). Sera were then two-fold serially diluted in PBS with and mixed 1:1 with four hemagglutinating units of H1N1 or H7N9 in 96 wells plates (starting dilution = 1:10). The serum-virus mixture was incubated for 20 minutes at RT, followed by the addition of 0.5% turkey red blood cells (bioTRADING) in a 1:1 mixture. Samples were incubated for 45 minutes at RT after which agglutination was scored.

### Virus neutralization assay

Virus neutralizing (VN) titers were determined as described previously [73] and according to WHO guidelines [72]. Sera were inactivated (30 minutes at 56 °C) and two-fold serially diluted in virus growth medium using a starting dilution of 1:8. Virus at a concentration of 100 TCID_50_ was added and the mixture was incubated for 2 hours at 37 °C. Next, the virus-serum mixture was transferred to 96 wells plates containing confluent MDCK cells and incubated for another 2 hours at 37 °C after which the medium was refreshed. Plates were incubated until a back titration plate reached CPE at a titer of 100 TCID_50_ (4-5 days). The 50% virus neutralization titers per ml serum was calculated by the Reed and Muench method [74].

### Pathology

Tissues harvested for histological examination (trachea, bronchus and left lung) were fixed in 10% neutral-buffered formalin, embedded in paraffin, sectioned at 4 *μ*m and stained with hematoxylin and eosin (HE) for examination by light microscopy. Semiquantitative assessment of influenza virus-associated inflammation in the lung (four slides with longitudinal section or cross-section of cranial or caudal lobes per animal) was performed on every slide as reported earlier [75] with few modifications: for the extent of alveolitis and alveolar damage we used: 0, 0%; 1, 1–25%; 2, 25–50%; 3, >50%. For the severity of alveolitis, bronchiolitis, bronchitis, and tracheitis we scored: 0, no inflammatory cells; 1, few inflammatory cells; 2, moderate numbers of inflammatory cells; 3, many inflammatory cells. For the presence of alveolar edema and type II pneumocyte hyperplasia we scored: 0, 0%, 1, <25%, 2, 25-50%, 3, >50%. The presence of alveolar hemorrhage we scored: 0, no; 1, yes. For the extent of peribronchial/perivascular edema we scored: 0, no, 1, yes. Finally, for the extent of peribronchial, peribronchiolar, and perivascular infiltrates we scored: 0, none; 1, one to two cells thick; 2, three to ten cells thick; 3, more than ten cells thick. Slides were examined without knowledge of the treatment allocation of the animals.

### Body temperature, body weight and lung weight

Temperature data were retrieved from the implanted temperature loggers and consisted of measurements taken every 30 minutes. Baseline temperature was calculated as the average temperature in the 5 days before infection. The change in temperature was calculated as deviation from baseline (ΔT). The area under the curve (AUC) was calculated as the total ΔT up till 5 dpi. Values smaller than ‘baseline - 2*standard deviation of baseline’ were excluded as these often occur due to anesthesia. Relative bodyweight and relative lung weight are expressed as a percentage of bodyweight or ratio on the day of infection.

### Data analysis

All the statistical tests carried out aimed at detecting differences between the distributions of responses in two treatment groups (e.g. H1N1/mRNA and placebo), each response pertaining to a given stimulus (or measured variable, e.g. body weight) on a given tissue on a given day. The tests are based on the ‘sum statistic’ [76] as implemented in the R package ‘coin’ [77], in the guise of the function ‘independence_test’, possibly with blocking in the event that some experiments were done on different days (in which case the data from the same experiment are collected in the same block), and with the (exact) p-values estimated by random permutations. The tests were grouped into various themes based on tissue and assay (e.g. all stimulations for lung IFNγ ELISpot), and the Benjamini-Hochberg (BH) method [78] was used separately per theme to control the false discovery rate (FDR) at the level of 10%. Only the results of the tests that passed through the BH method are reported and commented upon in the results section. The overall proportion of spurious results (over all the themes) is expected to be at most 10% of all those reported. Tables with the complete results of the tests and multiple testing corrections are available as Supplementary data file 1. The results reported are illustrated by graphs (e.g. box plots) in the main text or in the online supplemental material.

IFNγ-ELISpot spot counts, viral titers, serum titers and cell counts were log-transformed for statistical testing. We excluded two datapoints of flow cytometry data from data visualization and analysis. These datapoints (one in PBMC, one in lung) refer to the percentage IFNy^+^ within CD4^+^ T cells and were at least two-times higher than the nearest datapoint. No other data was excluded from analysis.

