## Supplemental Figures for "A universal influenza mRNA vaccine candidate boosts T-cell responses and reduces zoonotic influenza virus disease in ferrets"

**Abstract**

Universal influenza vaccines have the potential to protect against continuously evolving and newly emerging influenza viruses. T cells may be an essential target of such vaccines as they can clear infected cells through recognition of conserved influenza virus epitopes. We evaluated a novel T cell-inducing nucleoside-modified mRNA vaccine that encodes the conserved nucleoprotein, matrix protein 1 and polymerase basic protein 1 of an H1N1 influenza virus. To mimic the human situation, we applied the mRNA vaccine as a prime-boost regimen in naïve ferrets (mimicking young children) and as a booster in influenza-experienced ferrets (mimicking adults). The vaccine induced and boosted broadly-reactive T cells in the circulation, bone marrow and respiratory tract. Booster vaccination enhanced protection against heterosubtypic infection with potential pandemic H7N9 influenza virus in influenza-experienced ferrets. Our findings show that mRNA vaccines encoding internal influenza virus proteins are a promising strategy to induce broadly-protective T-cell immunity against influenza viruses.

### Supplemental figures

*Supplemental Table 1: Percentage amino acid identity of several proteins between influenza strains*

| Influenza virus | HA | NA | NP | M1 | PB1 |
|---|---|---|---|---|---|
| **H1N1** (A/California/07/2009) | 100 | 100 | 100 | 100 | 100 |
| **H2N2** (A/Leningrad/134/17/57) | 63,43 | 43,49 | 91,77 | 94,05 | 96,3 |
| **H3N2** (A/Uruguay/716/2007) | 43,01 | 44,03 | 89,56 | 92,46 | 97,49 |
| **H5N1** (A/Vietnam/1204/2004) | 62,9 | 84,22 | 93,78 | 95,63 | 96,43 |
| **H7N9** (A/Anhui/1/2013) | 41,12 | 45 | 92,97 | 92,46 | 95,77 |

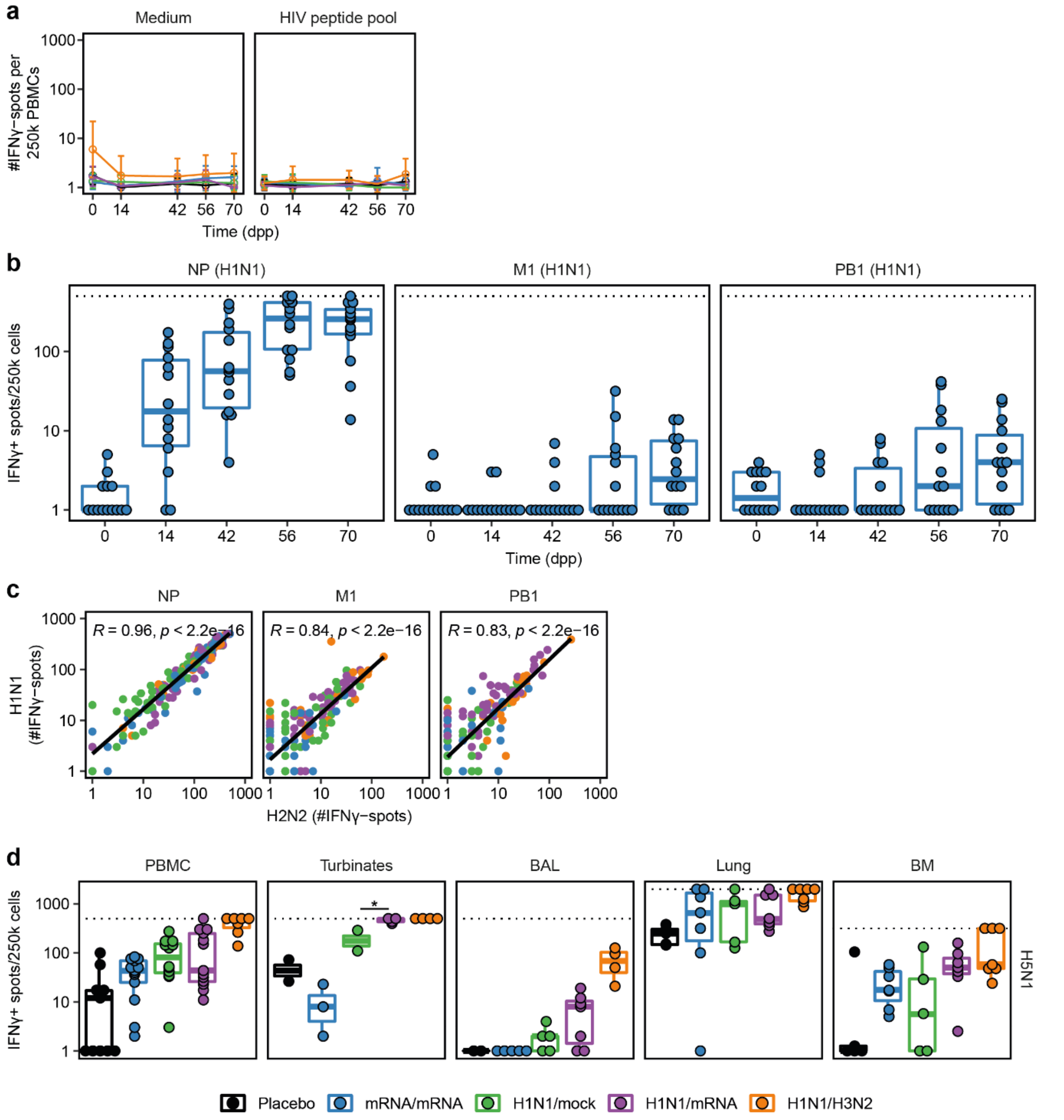


**Supplemental Figure 1:** Cellular responses measured by IFNγ ELISpot. **a)** Control stimulations for PBMC ELISpots performed 0, 14, 42, 56 and 70 days post priming (dpp). HIV responses were corrected for medium background responses. **b)** IFNγ responses in ferrets from mRNA/mRNA ferrets in time, showing that approximately half of the animals developed responses against M1 and PB1 after booster vaccination on 42 dpp. **c)** Correlation between peptide pool stimulations with NP, M1 and PB1 peptide pools of H1N1 (A/California/07/2009) and H2N2 (A/Leningrad/134/17/1957) influenza viruses. Data from ELISpots assays on 14, 42, 56 and 70 dpp are shown and correlation analysis was performed using the Pearson correlation coefficient (R). **d)** IFNγ responses measured by ELISpot after stimulation with H5N1 (A/Vietnam/1204/2004) influenza virus in various tissues 70 dpp. In panels b-d, each dot represents one animal, with n = 7-14 (PBMC), 2-4 (nasal turbinates) or 5-7 (BAL, lung, BM). For visualization purposes, only comparisons between groups mRNA/mRNA, H1N1/mock and H1N1/mRNA are shown. An overview of all statistical comparisons is detailed in Supplemental data file 1. * = p < 0.05, ** = p < 0.01, *** = p < 0.001.

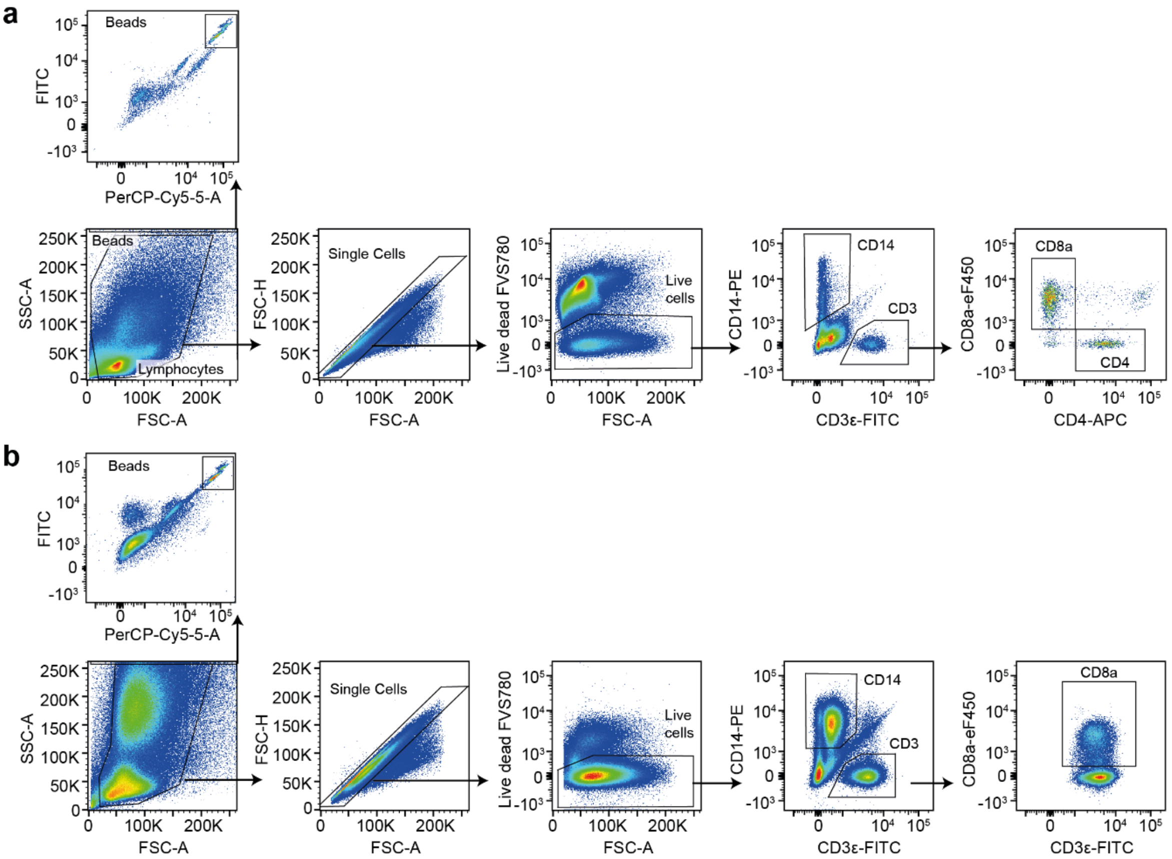


**Supplemental Figure 2:** FACS gating for cell counts in nasal turbinates and BAL. **a, b)** Gating strategy for cell populations in **a)** nasal turbinates and **b)** bronchoalveolar lavage (BAL). αCD4-APC staining was not consistent between BAL samples and was therefore excluded.

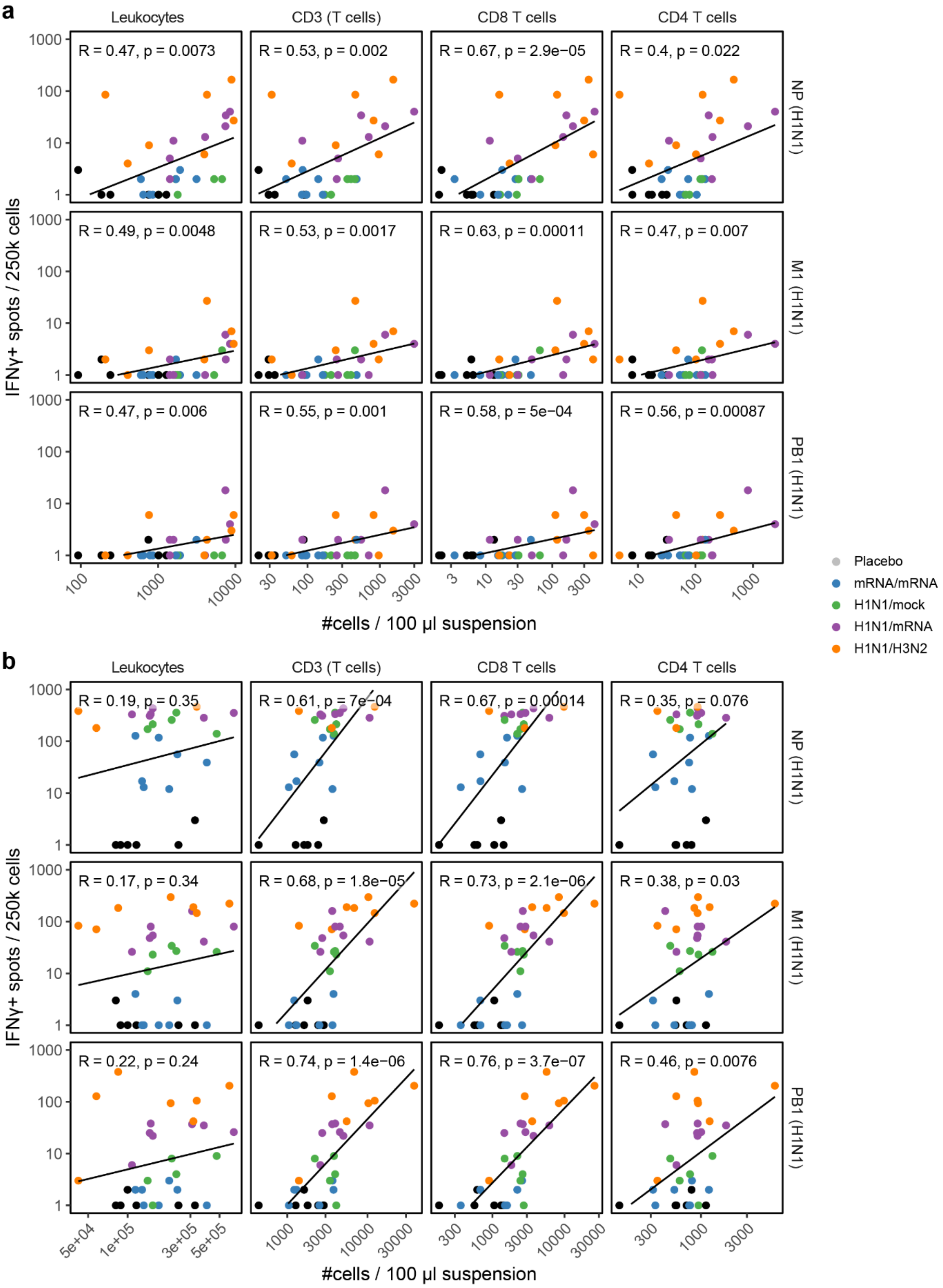


**Supplementary Figure 3:** Correlation between influx of cell populations and IFNγ spots in **a)** BAL fluid or **b)** nasal turbinates. Cell counts of different populations are plotted on the x-axis (data from Fig. 2c). IFNγ responses after H1N1 peptide pool stimulation are plotted on the y-axis (data from Fig. 2a). The correlation between cell counts and IFNγ responses is depicted by the black line and the correlation was calculated using the Pearson correlation coefficient (R). Each dot represents one animal with n = 5-7 per group.

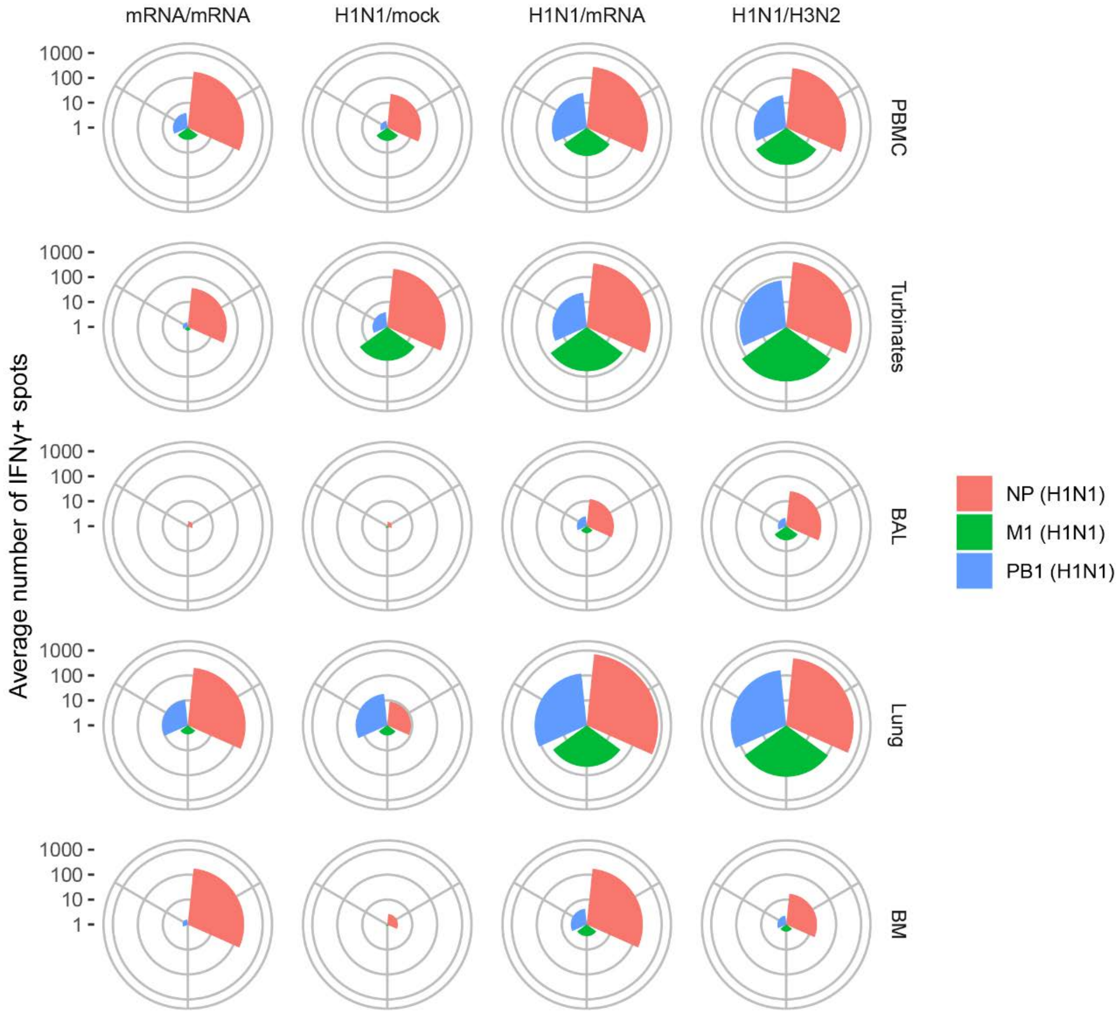


**Supplemental Figure 4:** Average cellular responses against NP, M1 and PB1 peptide pools of H1N1 influenza in different compartments at 70 days post priming. IFNγ ELISpot responses as depicted in Figure 1a and 2b,d,e were log-transformed and averaged per group for each response and tissue. Log-transformed means were transformed back to #spots per 250k cells and plotted per group and tissue. The size of each segment indicate the size of the average IFNγ response per group. For PBMCs, n = 12-14 for all groups except H1N1/H3N2 (n = 7). For all other tissues, n = 5-7.

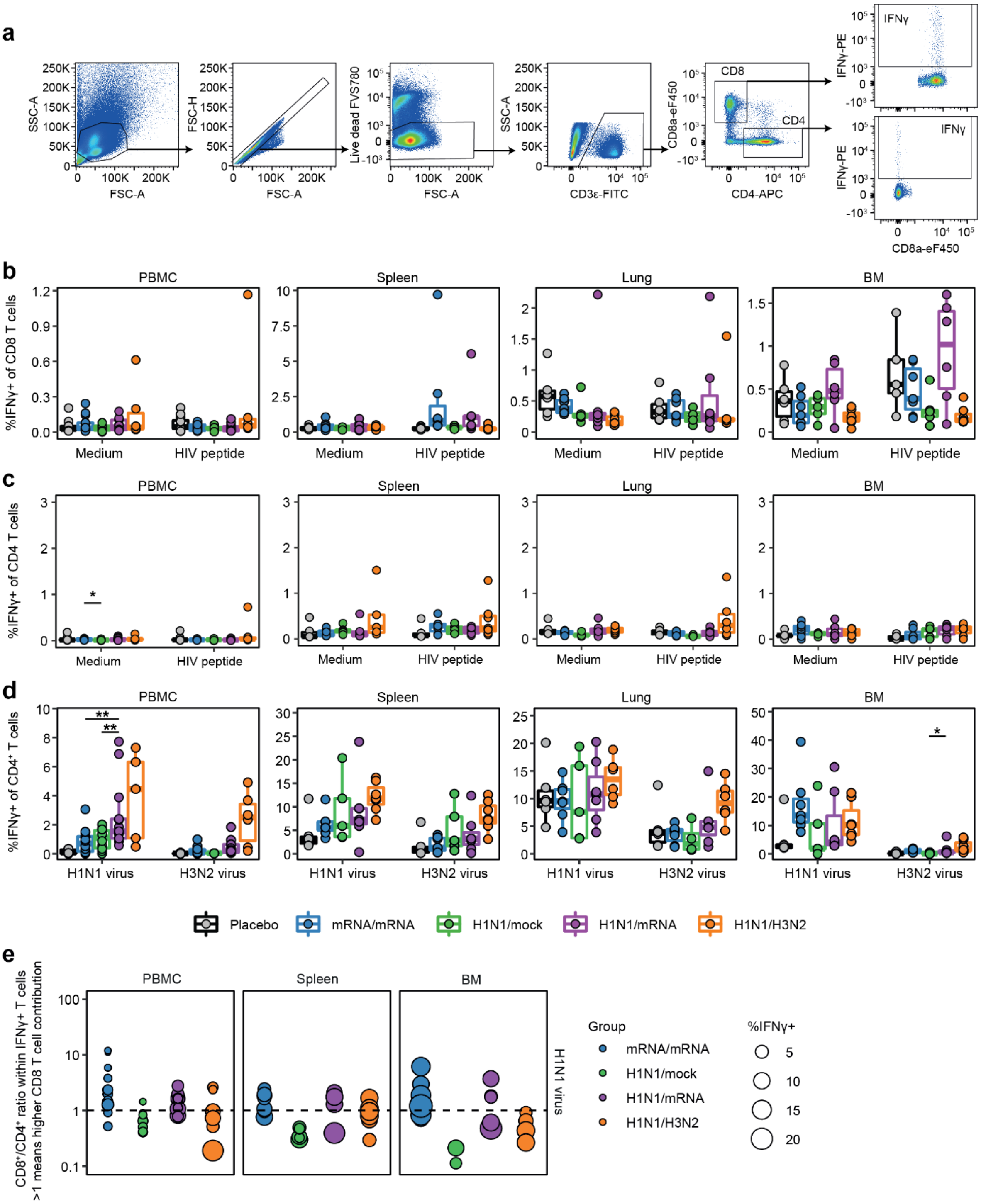


**Supplemental Figure 5:** IFNγ responses in $CD8^+$ and $CD4^+$ T cells measured by flow cytometry. **a)** Gating strategy for $IFN\gamma^+$ cells within the $CD8^+$ and $CD4^+$ T cell populations. Example shown is from PBMCs of an H1N1/mRNA-treated ferret after peptide cocktail stimulation (NP, M1 and PB1 peptide pools), but is representative for other tissues and groups. **b, c)** Negative controls for stimulations depicted in Figure 3. IFNγ expression in **b)** $CD8^+$ or **c)** $CD4^+$ T cells after stimulation of lymphocytes derived from blood, spleen, lung and BM with medium or HIV peptide pool. **d)** Percentage IFNγ-positive $CD4^+$ T cells after stimulation with H1N1 (A/California/07/2009) or H3N2 (A/Uruguay/217/2007) influenza viruses. **e)** Ratio between $CD8^+$ and $CD4^+$ T cells within the $CD3^+$ $IFN\gamma^+$ T-cell population after H1N1 influenza virus stimulation. Dotted line represents a ratio of 1 and samples with less than 50 $CD3^+$ $IFN\gamma^+$ cells were excluded from the analysis. No ratio was calculated for lung after H1N1 virus stimulation as high background responses were present in the $CD4^+$ population of placebo animals. Boxplots depict the median, 25% and 75% percentile, where the upper and lower whiskers extend to the smallest and largest value respectively within 1.5* the inter quartile ranges. In panels b-f, each dot represents one animal. In panels b-d, for PBMC n = 6-13 and for lung, BM and spleen n = 4-7. For visualization purposes, only comparisons between groups mRNA/mRNA, H1N1/mock and H1N1/mRNA are shown. An overview of all statistical comparisons is detailed in Supplemental data file 1. * = $p < 0.05$, ** = $p < 0.01$, *** = $p < 0.001$.

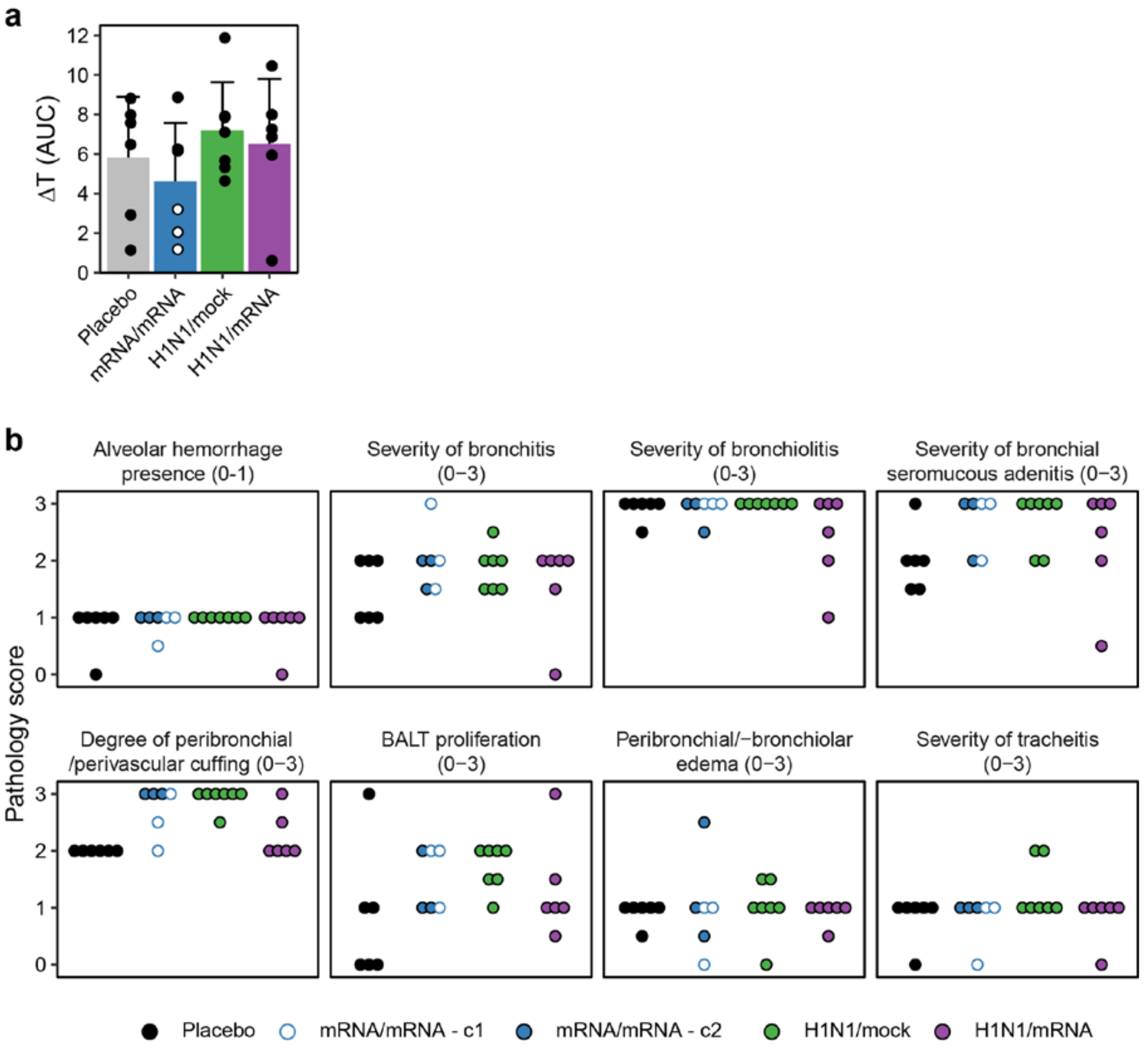


**Supplemental Figure 6:** Extended pathology during H7N9 influenza virus infection. **a)** Area under the curve (AUC) of body temperature, which was calculated from 0 to 5 days post infection (dpi) based on the data depicted in Fig. 4c. Values lower than mean + 2x SD were excluded as these are often due to anaesthesia. For one placebo ferret and three mRNA/mRNA ferrets (latter depicted in open circles), the AUC was only calculated up to 4 dpi due to these animals reaching the humane endpoints. Samples are depicted as mean ± SD with individual datapoints. **b)** Pathology scoring for various parameters. Each dot represents one animal. N = 6-7 for all panels. For visualization purposes, only comparisons between groups placebo, H1N1/mock and H1N1/mRNA are shown. No statistical testing was performed for panel b as these are nominal data. An overview of all statistical comparisons is detailed in Supplemental data file 1. * = p < 0.05, ** = p < 0.01, *** = p < 0.001.

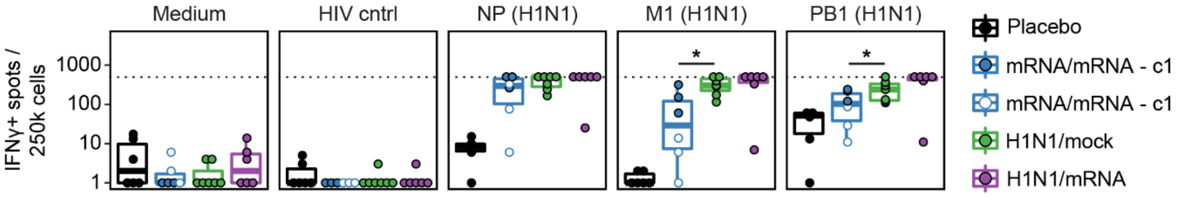


**Supplemental Figure 7:** Cellular responses during H7N9 influenza virus infection. PBMCs isolated 4 or 5 days post infection (dpi) with H7N9 (A/Anhui/1/2013) influenza virus were stimulated with medium, a HIV control peptide pool, or H1N1 peptide pools (NP, M1, PB1) in an IFNγ ELISpot assay. For one placebo ferret and three mRNA/mRNA ferrets, PBMCs were collected 4 dpi because these animals reached the humane endpoints and were sacrificed. For all other ferrets, blood was collected 5 dpi. Responses were corrected for medium background and data of 4 and 5 dpi were analysed together. Boxplots depict the median, 25% and 75% percentile, where the upper and lower whiskers extend to the smallest and largest value respectively within 1.5* the inter quartile ranges. Each dot represents one animal. In all panels, n = 6-7. For visualization purposes, only comparisons between groups placebo, H1N1/mock and H1N1/mRNA are shown. An overview of all statistical comparisons is detailed in Supplemental data file 1. * = $p < 0.05$, ** = $p < 0.01$, *** = $p < 0.001$.
